# Histone deacetylase 1 maintains lineage integrity through histone acetylome refinement during early embryogenesis

**DOI:** 10.1101/2022.05.05.490762

**Authors:** Jeff Jiajing Zhou, Jin Sun Cho, Han Han, Ira L. Blitz, Wenqi Wang, Ken W.Y. Cho

## Abstract

Histone acetylation is a pivotal epigenetic modification that controls chromatin structure and regulates gene expression. It plays an essential role in modulating zygotic transcription and cell lineage specification of developing embryos. While the outcomes of many inductive signals have been described to require enzymatic activities of histone acetyltransferases and deacetylases (HDACs), the mechanisms by which HDACs confine the utilization of the zygotic genome remain to be elucidated. Here, we show that histone deacetylase 1 (Hdac1) progressively binds to the zygotic genome from mid blastula and onward. The recruitment of Hdac1 to the genome at blastula is instructed maternally. *Cis*-regulatory modules (CRMs) bound by Hdac1 possess epigenetic signatures underlying distinct functions. We highlight a dual function model of Hdac1 where Hdac1 not only represses gene expression by sustaining a histone hypoacetylation state on inactive chromatin, but also maintains gene expression through participating in dynamic histone acetylation-deacetylation cycles on active chromatin. As a result, Hdac1 maintains differential histone acetylation states of bound CRMs between different germ layers and reinforces the transcriptional program underlying cell lineage identities, both in time and space. Taken together, our study reveals a comprehensive role for Hdac1 during early vertebrate embryogenesis.

## Introduction

A fundamental question in early development is the mechanism of zygotic genome activation (ZGA), which requires the degradation of maternal mRNAs and the activation of embryonic transcription (Tadros and Lipshitz, 2009). During ZGA, the embryonic genome undergoes a dramatic reprogramming of gene expression, which is also accompanied by remodeling of the embryonic epigenome. Post-translational modifications to histones are a major epigenetic regulation influencing chromatin structure and thus play a central role in ZGA. Histone acetylation appears during the onset of both minor and major ZGA waves in many species. In *Drosophila*, histone acetylation occurs at mitotic cycle 8 on a few early zygotic genes (Li et al., 2014). miR430, the first zygotically active gene, is marked by H3K27ac in 64-cell staged zebrafish embryos (Chan et al., 2019). Genome-wide H3K27ac is detected at mid blastula shortly after the onset of ZGA in *Xenopus* (Gupta et al., 2014). In mice, the zygotic genome is increasingly marked by H3K27ac from immature and metaphase II oocytes to 2-cell-stage embryos (Dahl et al., 2016). Despite these findings, there remain several major questions. How is the interplay of enzymes regulating histone acetylation employed in developing embryos? What is the role of observed histone acetylation on gene expression? How are the spatial and temporal patterns of histone acetylation established during ZGA?

Histone acetylation occurs on the ε-amino group of the lysine residues within *N*-terminal tails of all four core histones (Inoue and Fujimoto, 1969; Seto and Yoshida, 2014). Acetylation is a reversible process that is directly catalyzed by opposing activities of histone acetyltransferases (HATs) and histone deacetylases (HDACs). In addition, HATs and HDACs can also regulate the acetylation of lysine residues on non-histone proteins (Choudhary et al., 2009). Histone acetylation is often associated with active gene transcription because the acyl groups neutralize the positive charge on the lysine residues, thereby reducing the affinity of histones to DNA (Wang et al., 2000; Anderson et al., 2001); it also serves as a binding platform for bromodomain (BRD) proteins which scaffold and stimulate the transcriptional machinery (Hassan et al., 2007; Filippakopoulos et al., 2012). The balance between HATs and HDACs directly shapes histone acetylation landscapes and subsequently affects transcriptomes.

HDACs are critical epigenetic regulators because they reset chromatin states by returning acetylated lysine residues on histones to the basal state, which can subsequently be subjected to alternative modifications such as methylation. HDACs are grouped into four classes based on phylogenetic conservation. Class I (HDAC1, 2, 3, 8), Class II (HDAC4, 5, 6, 7, 9, 10), and Class IV (HDAC11) HDACs are zinc-dependent and are related to yeast Rpd3, Had1, and Hos3 respectively; Class III (SIRT1, 2, 3, 4, 5, 6, 7) HDACs, also known as Sirtuins, are NAD+-dependent and are related to yeast Sir2 (Gregoretti et al., 2004; Milazzo et al., 2020). HDACs are well-characterized negative regulators of gene expression during development. For example, HDAC1 silences homeotic genes in cooperation with Polycomb group repressors in *Drosophila* (Chang et al., 2001). In zebrafish, Hdac1 represses Notch targets during neurogenesis (Cunliffe, 2004; Yamaguchi et al., 2005). In *Xenopus*, HDAC activity suppresses Vegt-induced ectopic mesoderm in ectoderm lineages (Gao et al., 2016), represses multi-lineage marker genes at blastula (Rao and LaBonne, 2018) and desensitizes dorsal Wnt signaling at late blastula (Esmaeili et al., 2020). Conversely, HDACs can also positively regulate gene expression. For instance, inhibition of HDAC activities rapidly down-regulates some genes in yeast, suggesting an activator function of HDACs (Bernstein et al., 2000). Genetic deletions or pharmacological application of HDAC inhibitors in cell lines results in both up- and down-regulated genes (Reid et al., 2005; Zupkovitz et al., 2006; Meganathan et al., 2015). Furthermore, genome-wide studies showed that HDACs occupy genomic loci of active genes, and their binding correlates with gene activities (Kurdistani et al., 2002; Wang et al., 2002; Wang et al., 2009; Kidder et al., 2012). These seemingly opposing functions of HDACs raise an important question as to the exact roles of HDACs on chromatin states and transcriptomes in developing embryos.

In this study, we focus on the role of Hdac1 in regulating the zygotic epigenome and transcriptome during *Xenopus* germ layer specification coinciding with ZGA. Current evidence in *Xenopus* as well as in other non-mammalian systems suggests that the early embryonic genome is rather naïve and that major chromatin modifications occur at or after ZGA (Bonn et al., 2012; Vastenhouw et al., 2010; Gupta et al., 2014; van Heeringen et al., 2014). Thus, the system allows us to probe the earliest establishment of histone acetylation and dissect the link between actions of Hdac1, the zygotic histone acetylome, and zygotic transcriptome during the first cell lineage segregation event. Here, we report that the major Hdac1 binding to the embryonic genome occurs at blastula; the binding of Hdac1 during this stage is dependent on maternal factors. We highlight a dual function model for Hdac1. First, Hdac1 keeps inactive chromatin free of histone acetylation, preventing gene misactivation in respective germ layers. Second, Hdac1 participates in dynamic histone acetylation-deacetylation cycles on active chromatin, sustaining the expression of genes that are enriched in respective germ layers. Taken together, our study reveals a coordinated spatial and temporal regulation by Hdac1 during ZGA.

## Results

### Hdac1 binds to the genome progressively during blastula and onward

In order to identify the major functional candidates of HDACs during the early *Xenopus* embryogenesis, we examined the temporal RNA expression profiles (Owen et al., 2016) of HDAC family members (Class I, Class II, and Class III HDACs) from the zygote to the beginning of the neurula stage in *Xenopus tropicalis*. The RNA expression level of *hdac1* is the highest among all HDACs examined, followed by *hdac2* (Figure S1A). Both Hdac1 and Hdac2 proteins are present in the unfertilized egg to the mid gastrula stage and the overall expression levels of Hdac1 and Hdac2 are relatively constant during this period (Figure 1A). These data reveal that Hdac1/2 are the major maternally endowed HDACs functioning during this window of development.

**Figure 1.**
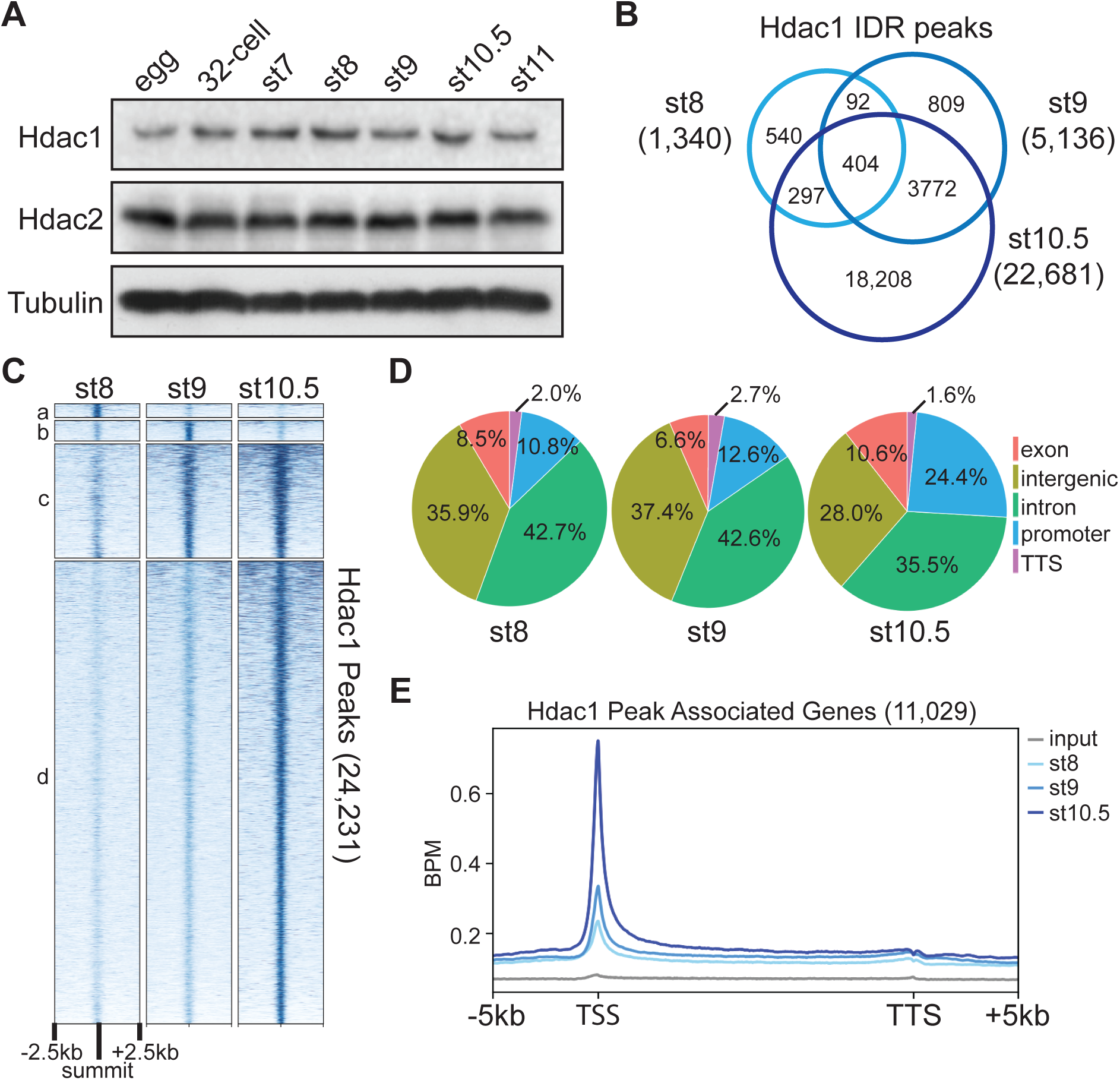
Hdac1 binds to genome gradually during early *Xenopus* development. (**A**) Western blot analyses showing Hdac1 and Hdac2 proteins over a time course of early development. *β*-tubulin is used as a loading control. (**B**) Venn diagram comparing Hdac1 IDR peaks among examined stages. The sum of peaks in st8 and st9 are smaller due to instances where more than one peak from st8 or st9 overlaps the single same st10.5 peak. (**C**) Clustered heatmap showing Hdac1 ChIP-seq signals at each stage over a window of 5kb centered on the summit of all Hdac1 IDR peaks in descending order. (**D**) Distributions of Hdac1 IDR peaks at each stage across five defined genomic features. The promoter is defined as −1kb to +100bp from TSS (transcription start site) while the TTS (transcription termination site) is defined as −100bp to +1kb from TTS. (**E**) Distributions of Hdac1 ChIP-seq signals within the intervals of 5 kb upstream of gene model 5’ ends, gene bodies (normalized for length), and 5 kb downstream of gene model 3’ ends at each stage. The signal of st9 input DNA ChIP-seq is used as a negative control. Y-axis values represent reads quantified by bins per million (BPM) at a bin size of 50 bp.

Since Hdac1 modulates various aspects of transcriptional regulation and the chromatin landscape, we examined genome-wide Hdac1 binding patterns during the early germ layer development using chromatin immunoprecipitation assays followed by sequencing (ChIP-seq) (Figure S1B). A set of high-confidence peaks at each stage were obtained using irreproducibility discovery rate (IDR) analysis (Li et al., 2011) from two biologically independent samples (Figure S1C). IDR analysis reveals that Hdac1 binds to 1,340 regions at the mid blastula (st8), 5,136 regions at the late blastula (st9), and 22,681 regions at the early gastrula (st10.5) stages (Figure 1B). Overall, a minority of Hdac1 peaks are unique to each of the blastula stages (Cluster a and b) while a majority of Hdac1 peaks are present across multiple stages (Cluster c) and at the early gastrula stage (Cluster d) (Figure 1C, S1D). These results reveal that Hdac1 is progressively directed to the genome during early development.

To investigate the differences of Hdac1 genomic occupancy across early stages, we examined various genomic features of regions bound by Hdac1 at each stage. The majority of Hdac1 peaks are found in either intergenic or intronic regions; a minor fraction of Hdac1 peaks is present at exons or transcriptional termination sites. A notable observation is the increased Hdac1 binding to an increased amount of promoter regions over developmental times (Figure 1D). We then analyzed the distribution of Hdac1 across 3 stages along the gene bodies of genes bound by Hdac1. A higher enrichment of Hdac1 binding is located near the promoters of genes as development proceeds (Figure 1E). The increased binding of Hdac1 around promoters infers that Hdac1 is involved in the epigenetic regulation of these genes during ZGA.

### Blastula Hdac1 binding is maternally instructed

Since Hdac1 binds progressively to the genome during ZGA, we investigated the contribution of zygotic factors to the recruitment of Hdac1 at the late blastula stage. We injected α-amanitin, which blocks both transcription initiation and elongation (Chafin et al., 1995), to block embryonic transcription (Figure S2A). A high Pearson correlation (0.93) between α-amanitin-injected and control embryos (CT) is observed at all Hdac1 IDR peaks (Figure 2A), which is higher than the Pearson correlation (0.87) representing the batch effect when CT is compared to a different batch of same staged embryos (WT). There are no significant signal differences between α-amanitin-injected and control embryos at all Hdac1 IDR peaks (Figure S2B). We conclude that Hdac1 binding is independent of zygotic transcription at the late blastula stage, suggesting the importance of maternal factors in Hdac1 recruitment.

**Figure 2.**
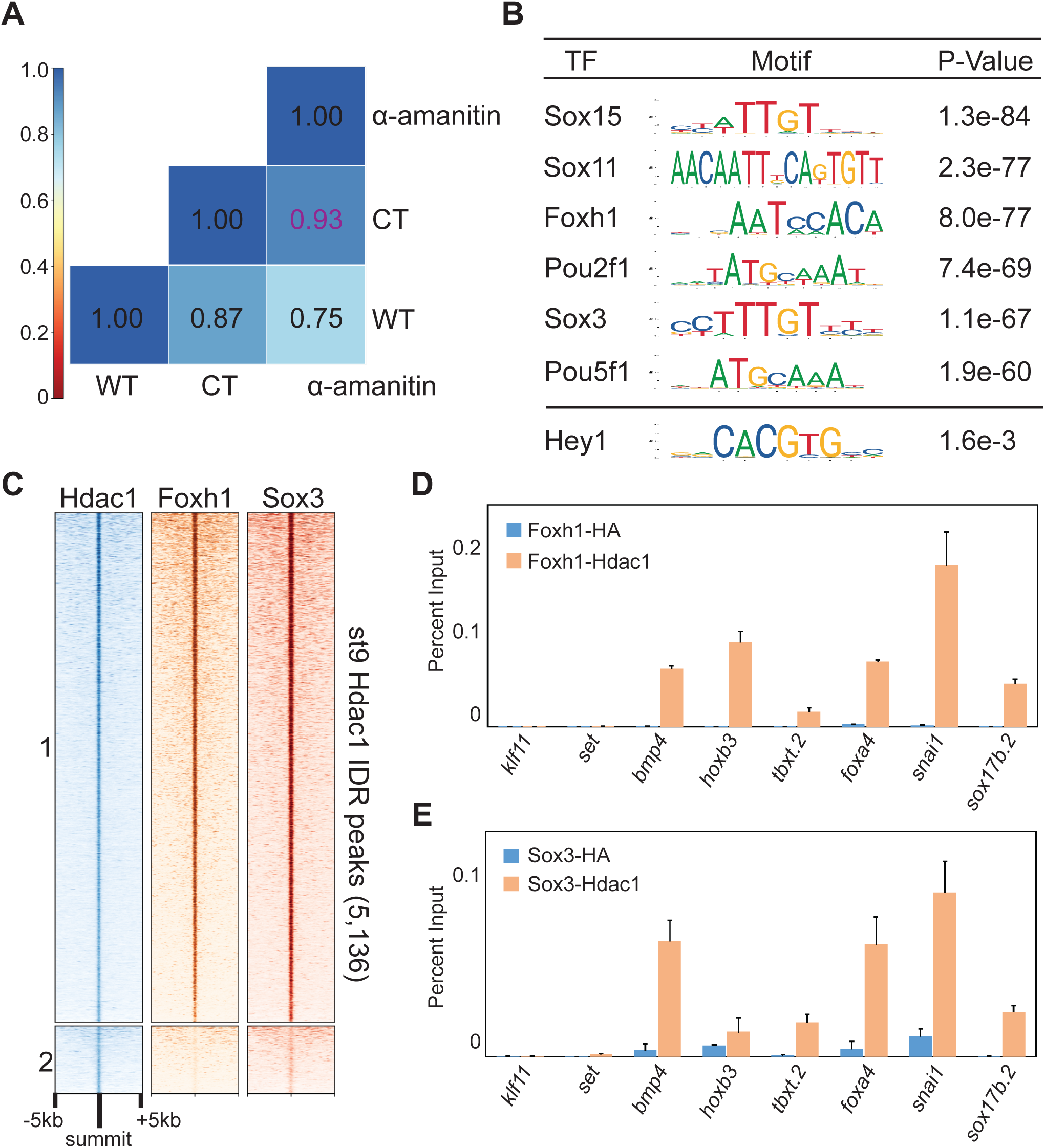
Maternal factors instruct Hdac1 recruitment during blastula stages. (**A**) Pairwise Pearson correlation analyses comparing ChIP-seq signals of st9 Hdac1 IDR peaks between listed samples. WT denotes samples from time-course experiments; α-Amanitin denotes α-amanitin injected embryos and CT denotes batch-matched control uninjected embryos. (**B**) Motif analyses of st9 Hdac1 peaks (500bp centered on IDR peak summit). Motif sequence to the corresponding factor is retrieved from JASPAR. Hey1 is an example of TF motif with low significance. (**C**) Clustered heatmap depicting st9 Foxh1 and Sox3 ChIP-seq signals in a window of 10kb centered on st9 Hdac1 IDR peaks with descending order. (**D-E**) St9 sequential ChIP-qPCR analyses for (**D**) Foxh1 and Hdac1 co-bound regions and (**E**) Sox3 and Hdac1 co-bound regions. anti-HA is used as a negative control. The error bars represent the variation from two technical replicates.

To identify maternal factors that facilitate the recruitment of Hdac1 to the genome, we performed a *de novo* motif search of the DNA sequences of 5,136 st9 Hdac1peaks. Sox, Foxh1, and Pou motifs are found to be the most frequent maternal TF motifs (Figure 2B). We thus compared the genomic binding profiles of Hdac1 to two maternal TFs, Foxh1 (Charney et al., 2017) and Sox3. A majority of Hdac1 bound regions (Cluster 1) overlaps with both Foxh1 and Sox3 bound regions while only a small fraction of Hdac1 bound regions (Cluster 2) overlaps the binding of either Foxh1, or Sox3, or neither (Figure 2C). More than 80% of Hdac1 peaks overlap with Foxh1 or Sox3 peaks (Figure S2C). A positive correlation between Hdac1 binding and Foxh1/Sox3 binding is observed at all Hdac1 IDR peaks (Figure S2D). We noted frequent overlapping binding of Hdac1 with Foxh1/ Sox3 (Figure S2E), and highly enriched signals of Hdac1, Foxh1, and Sox3 present around promoters of genes (Figure S2F). All these observations suggest a role for Foxh1 and Sox3 in Hdac1 recruitment. We also confirmed the co-occupancy of Hdac1 with each of Foxh1 and Sox3 TFs on the same DNA molecules using sequential ChIP-qPCR (Figure 2D, 2E, S2G). Taken together, we propose that maternal TFs such as Foxh1 and Sox3 facilitate Hdac1 recruitment.

### Hdac1 binds to genomic regions with distinct epigenetic signatures

To further characterize regions bound by Hdac1 across early germ layer development, we examined epigenetic signatures (Gupta et al., 2014; Hontelez et al., 2015; Charney et al., 2017) on Hdac1 peaks across various stages. A high signal density of Ep300 (a HAT) is observed in Hdac1 peaks (Cluster b, c, and d) from the late blastula and onward where RNA polymerase II signals also emerge (Figure S3A). This indicates that Hdac1 and Ep300 share similar binding profiles on transcriptionally active genes. We next surveyed several histone acetylation modifications on regions bound by Hdac1 (Figure 3A). Consistent with the overlapping binding of Ep300, Hdac1 peaks display a moderate level of H3K9ac (Cluster c and d), high levels of H3K18ac (Cluster b, c, and d), H3K27ac (Cluster b, c, and d), and pan-H3 lysine acetylation (pan-H3Kac) (Cluster b, c, and d). We then examined several histone methylation modifications that are associated with gene activation (Figure S3B). H3K4me1, a primed enhancer mark (Creyghton et al., 2010), displays a moderate level of signal density at Hdac1 bound regions (Cluster c and d); H3K4me3, an active promoter mark (Heintzman et al., 2007), displays a high level of signal density at Hdac1 bound regions (Cluster c and d); H3K36me3, a transcription elongation mark (Kolasinska-Zwierz et al., 2009), displays minimal signal density at any Hdac1 bound regions. Taken together, these observations reveal that Hdac1 binds to genomic regions with active epigenetic signatures.

**Figure 3.**
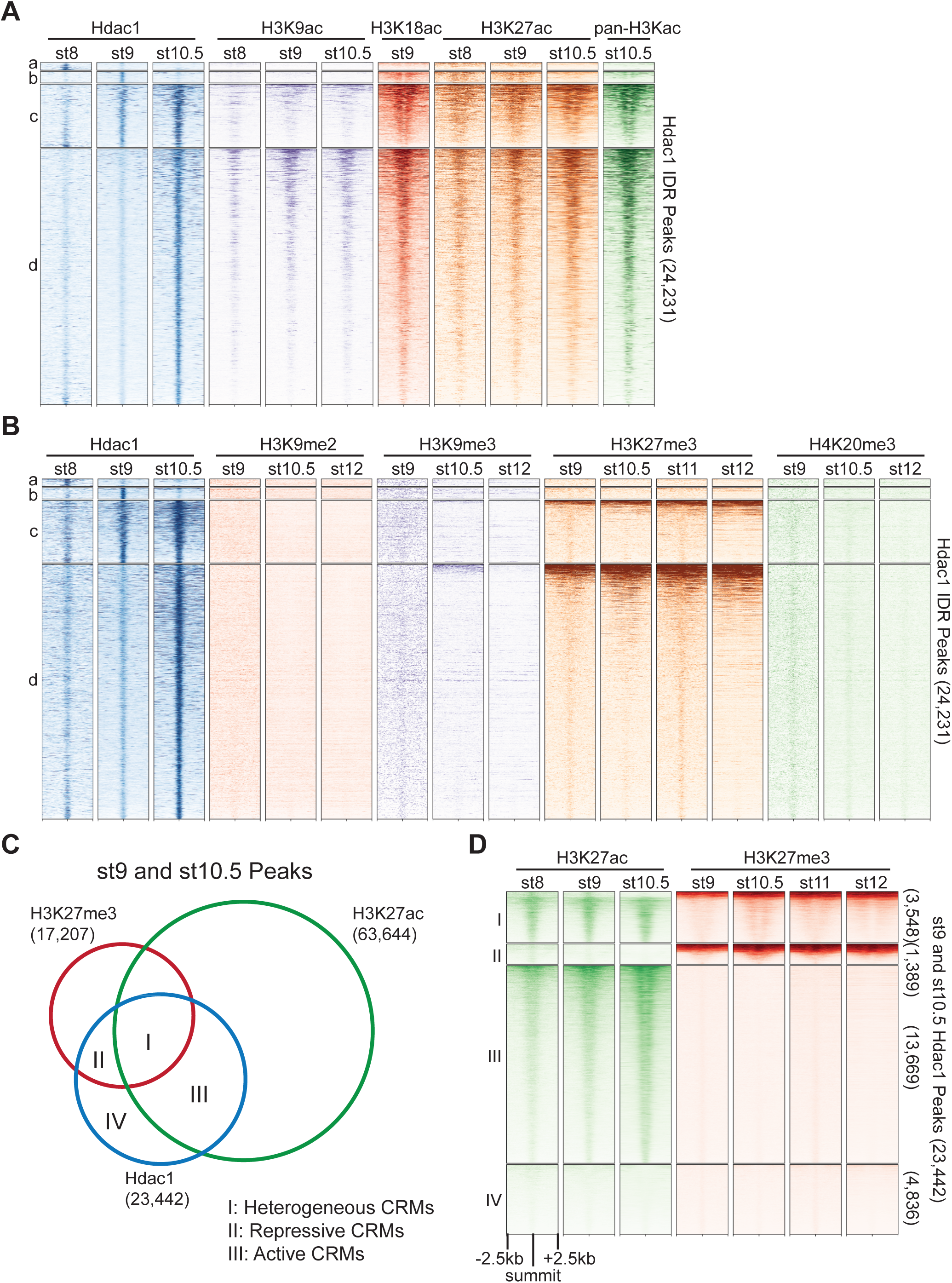
Hdac1 binds to CRMs containing functionally distinct histone modifications. (**A, B**) Clustered heatmaps showing signals from several stages of various (**A**) histone acetylation marks and (**B**) repressive histone methylation marks on Hdac1 peaks. Each cluster corresponds to the same regions present in Figure 1C. (**C**) Venn diagram illustrating Hdac1 peaks overlapping with H3K27me3 and H3K27ac peaks from both st9 and st10.5 combined. (**D**) Clustered heatmaps depicting signals of H3K27me3 and H3K27ac across multiple stages on st9 and st10.5 Hdac1 combined peaks. Clusters denote the same genomic regions in Venn diagram regions demarcated in Figure 3C. In panels A, B and D, numbers on the right side indicate the total number of regions in each cluster. The signals are shown in a window of 5kb centered on the summits of Hdac1 peaks presented in descending order of track signal intensities within each cluster.

Hdac1 resets the state of acetylated histones by removing acetyl groups, thus can facilitate the formation of repressive chromatin. Here we analyzed several histone methylation modifications associated with gene repression (Figure 3B). Hdac1 binds to genomic regions mostly free of H3K9me2, H3K9me3, or H4K20me3. All three modifications are known to mark constitutive heterochromatin denoting gene-poor areas consisting of tandem repeats (Richards and Elgin, 2002). In addition, a fraction of Hdac1 bound regions (Cluster c and d) display a strong signal density of H3K27me3 (Figure 3B), which is known to mark facultative heterochromatin consisting of developmental-cue silenced genes (Trojer and Reinberg, 2007). These observations suggest that Hdac1 binds to facultative heterochromatic regions facilitating the repression of genes instructed by developmental programs.

Since Hdac1 peaks (Cluster b, c, d, but not a) are marked by both active and repressive epigenetic signatures, we wonder whether Hdac1 functions differently at epigenetically distinct genomic loci. We compared peaks between two functionally opposing histone modifications H3K27ac and H3K27me3 to Hdac1 peaks, then subdivided these Hdac1 peaks into 4 clusters representing functionally distinct CRM types (Figure 3C, S3C). Cluster I denotes 3,548 Hdac1 peaks marked by both H3K27ac and H3K27me3 (Figure 3D). Given that both H3K27ac and H3K27me3 are subjected to the same lysine residue, we speculate that these regions are differentially marked in space due to heterogeneous cell populations present in the whole embryo. Hdac1 Cluster I peaks are referred to as heterogeneous CRMs. Cluster II denotes 1,389 Hdac1 peaks marked by only H3K27me3 indicating that these regions are associated with inactive developmental genes (Figure 3D). Hdac1 Cluster II peaks represent repressive CRMs. Cluster III denotes 13,669 Hdac1 peaks marked by only H3K27ac suggesting that these CRMs are associated with active genes (Figure 3D). Hdac1 Cluster III peaks are considered as active CRMs. Cluster IV denotes 4,836 Hdac1 peaks with neither H3K27ac nor H3K27me3 modifications (Figure 3D). Together, we show that different Hdac1 bound regions are subjected to functionally distinct epigenetic modifications suggesting that Hdac1 is differentially utilized at different CRMs.

### HDAC activity is required for differential germ layer histone acetylomes

The major function of HDACs is to catalyze the removal of acetyl groups from histones. We hypothesize that Hdac1 differentially regulates histone acetylation of four different Hdac1 CRM clusters (Clusters I-IV in Figure 3C). To test this hypothesis, we treated embryos continuously with a widely used pan-HDAC inhibitor, Trichostatin A (TSA) (Yoshida et al., 1990) beginning at the 4-cell stage and followed the development up to tailbud stages. Embryos treated with TSA are developmentally arrested at gastrula (Figure 4A). We observed the presence of dorsal blastopore lip suggesting that the progression but not the initiation of gastrulation is defective. A Class I HDAC inhibitor, Valproic acid (VPA) (Göttlicher et al., 2001), also produces a similar phenotype (Figure S4A). To investigate whether HDAC inhibition leads to changes in overall histone acetylome, we surveyed six well-known histone acetylation modifications between TSA- and solvent-treated embryos by western blot. Indeed, drastically increased levels of H3K9ac, H3K18ac, and H3K27ac are observed in TSA-treated embryos (Figure 4B; Rao and LaBonne, 2018) while H3K14ac, H3K56ac, and H4K20ac are not detected during this stage of development. These data indicate that HDAC activity is required to maintain the proper level of histone acetylation during gastrulation.

**Figure 4.**
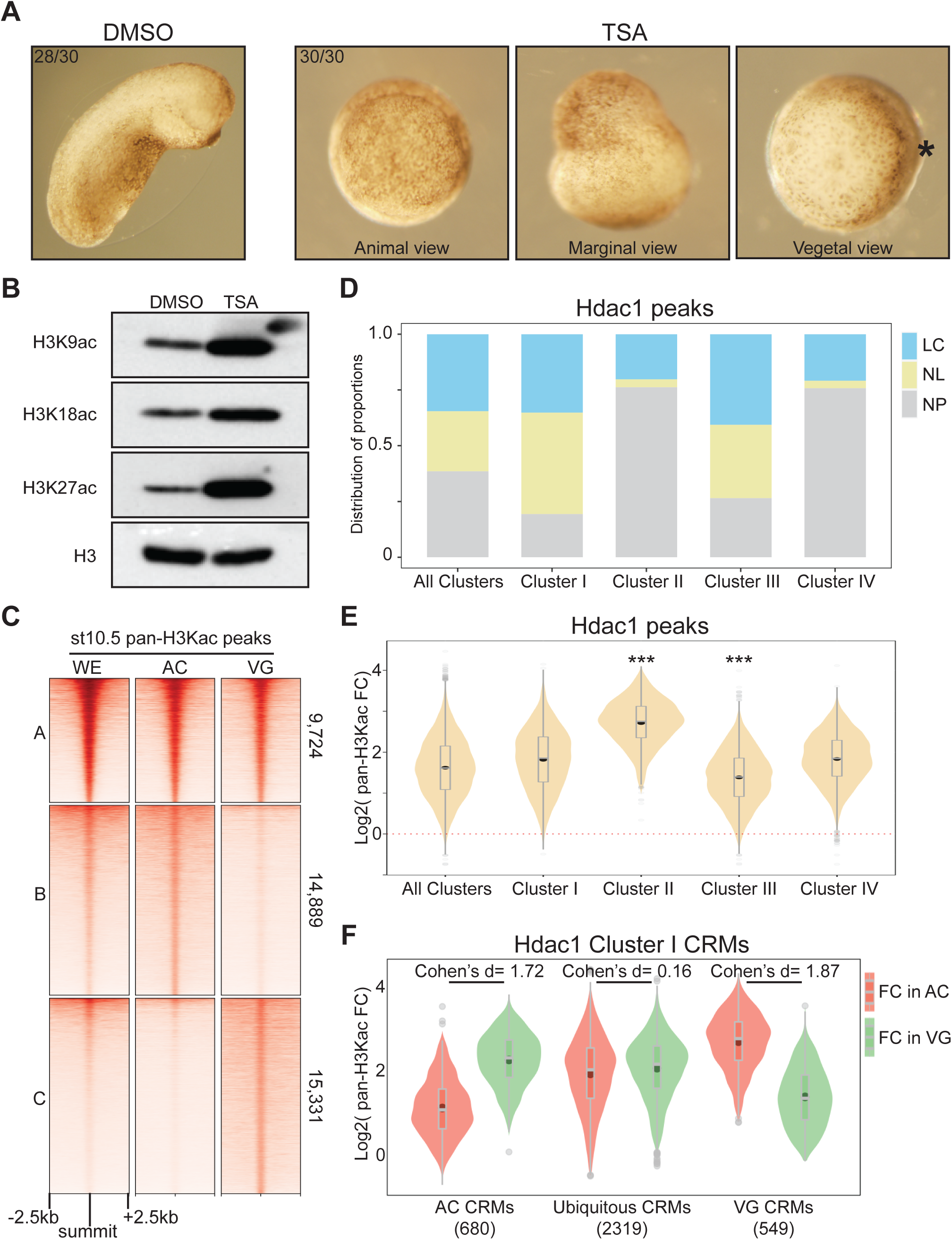
Hdac1 maintains differential H3 acetylomes between germ layers. (**A**) Embryos treated with 100uM TSA displaying gastrulation defects 24 hours post-fertilization. Asterisk denotes the dorsal side containing the early blastopore lip. (**B**) Western blot analyses showing various histone acetylation modifications affected by HDAC inhibition. anti-H3 is used as a loading control. (**C**) Clustered heatmap depicting signals of pan-H3Kac at st10.5 in whole embryos (WE), animal cap cells (AC), and vegetal mass cells (VG). The signals are shown in a window of 5kb centered on the summits of combined AC and VG peaks presented in descending order of track signal intensities within each cluster. (**D**) Stacked bar graph representing proportions of localized versus non-localized pan-H3Kac signals found at Hdac1 CRM clusters in Figure 3C. A Hdac1 peak is considered to exhibit localized pan-H3Kac if it overlaps with either AC- or VG-specific pan-H3Kac peaks (Cluster B and Cluster C in Figure 4C); a Hdac1 peak is considered to exhibit non-localized pan-H3Kac if it overlaps with pan-H3Kac peaks shared between AC and VG (Cluster A in Figure 4C). LC: localized; NL: non-localized; NP: not overlap with any pan-H3Kac peak (**E**) Fold changes (FC) of pan-H3Kac signals at Hdac1 CRM clusters (clusters in Figure 3C). Red dotted line denotes the level of zero. *** denotes *p*< 0.001 (Student’s *t*-test). (**F**) Fold changes (FC) of pan-H3Kac signals in Cluster I of Hdac1 CRM clusters (clusters in Figure 3C) categorized spatially (described in Supplemental Methods).

Given that Hdac1 CRM clusters (Clusters I-IV in Figure 3C) are subjected to both active and repressive epigenetic modifications presumably in different germ layers, we compared the general status of H3 acetylome (pan-H3Kac) between two distinct germ layers, the animal cap (AC, presumptive ectoderm) and the vegetal mass (VG, presumptive endoderm). We performed pan-H3Kac ChIP-seq at early gastrula (st10.5) on dissected AC and VG explants from embryos treated with either TSA or solvent control (Figure S4B). A comparison of high-confidence peaks from each explant reveals that a majority (∼ 75%) of pan-H3Kac are specifically marked in ectodermal (∼ 37%) and endodermal (∼ 38%) germ layers (Figure 4C, S4C). To correlate the differential pan-H3Kac states and the differential gene expression profiles between two germ layers, we assigned high enrichment pan-H3Kac peaks (∼ top 30%) to the nearest genes within 10kb and compared the expression levels of these genes in each germ layer (Blitz et al., 2017). Indeed, the expression level of genes enriched with AC-specific pan-H3Kac is higher in AC than VG (Figure S4D), and the expression level of genes enriched with VG-specific pan-H3Kac is higher in VG than AC (Figure S4E). Well-known genes with germ-layer specific expression exhibit localized pan-H3Kac signals between germ layers (Figure S4F). These results illustrate that the histone acetylation profile generally coincides with the animally and vegetally localized expression of transcripts.

To uncover the role of Hdac1 in regulating histone acetylation states of CRM clusters (Clusters I-IV in Figure 3C), we first examined the general distribution of pan-H3Kac in these Hdac1 CRM clusters. Consistent with H3K27 modifications, we observed that the majority of both heterogeneous CRMs (Cluster I, both H3K27ac and H3K27me3) and active CRMs (Cluster III, only H3K27ac), but not repressive CRMs (Cluster II, only H3K27me3) are marked by pan-H3Kac; near half of pan-H3Kac marks on heterogeneous or active CRMs are localized either animally or vegetally (Figure 4D). The localized pan-H3Kac marks on active CRMs suggests that these CRMs are regionally active in specific germ layers. We then quantitatively examined the levels of pan-H3Kac signals on each CRM within clusters I-IV (Figure 3C) after TSA treatment using a ChIP spike-in strategy (Egan et al., 2016). A global increase of pan-H3Kac signals across all Hdac1 CRM clusters is observed (Figure S4G), which is consistent with the result by western blot (Figure 4B). We found that CRMs of Hdac1 clusters I-IV respond differently upon HDAC inhibition: repressive CRMs (Cluster II, only H3K27me3) show a significantly higher acquisition (∼ 5-fold increase) of pan-H3Kac signals, while active CRMs (Cluster III, only H3K27ac) show the lowest acquisition (∼ 3-fold increase) of pan-H3Kac signals when compared to other clusters (Figure 4E). Collectively, these results reveal that repressive CRMs are prone to loss of HDAC activity resulting in histone hyperacetylation, while active CRMs gain a modest increase of histone acetylation upon HDAC inhibition.

Lastly, we explored how Hdac1 CRM clusters (Clusters I-IV in Figure 3C) differentially respond to HDAC inhibition in specific germ layers. For heterogeneous CRMs (Cluster I, both H3K27ac and H3K27me3), we found that the fold increases of pan-H3Kac signals after TSA treatment are negatively correlated with the levels of endogenous pan-H3Kac signals in both AC and VG explants (Figure S4H). Next, heterogeneous CRMs were subdivided into 3 spatial categories: pan-H3Kac enriched animally (AC CRMs), pan-H3Kac enriched vegetally (VG CRMs), and ubiquitous CRMs. We then calculated the fold changes of pan-H3Kac signals after TSA treatment for each CRM within each spatial category. Interestingly, germ-layer specific AC and VG CRMs have a much greater effect size (1.72 for AC CRMs and 1.87 for VG CRMs) when compared to ubiquitous CRMs (0.16) upon HDAC inhibition (Figure 4F). This indicates that AC CRMs acquire a drastic increase of pan-H3Kac signals in VG but not in AC in response to the loss of HDAC activity; likewise, VG CRMs acquire a drastic increase of pan-H3Kac signals in AC but not in VG upon the loss of HDAC activity. A similar trend is also observed on active CRMs (Cluster III, only H3K27ac) emphasizing germ layer-specific functions of these CRMs (Figure S4I, S4J). Taken together, these data demonstrate that Hdac1 maintains differential H3 acetylation states between germ layers through its catalytic activity.

### HDAC activity modulates developmental genes between germ layers

Histone deacetylation is generally associated with transcriptional repression; hence, HDACs are considered as transcriptional corepressors. To determine how Hdac1 CRM clusters I-III (Figure 3C) influence the activities of their corresponding genes, each CRM within a cluster was assigned to the nearest gene located within 10 kb and then subdivided these genes into 3 classes (Figure 5A). Class 1 denotes 3,104 genes that have CRMs with mixed marks of H3K27ac and/or H3K27me3 suggesting that these genes are differentially expressed in different germ layers. Class 2 denotes 629 genes whose CRMs are marked with only H3K27me3 indicating that these genes may be repressed. Class 3 denotes 5,913 genes whose CRMs are marked with only H3K27ac inferring that these genes are active in various germ layers. We first investigated the temporal expression pattern (Owens et al., 2016) of each gene class. To exclude the interference posed by residual maternal transcripts at these early stages, we examined the expression patterns of exclusively zygotic genes in each class and found that Class 1 and Class 3 genes are gradually activated after ZGA while Class 2 genes remain mostly silent even at late gastrula (Figure 5B). To assess whether Class 2 genes remain inactive throughout the development, we extended our temporal expression analysis through to tailbud stage 26 (23 hpf). Class 1 and Class 3 genes are continuously active after ZGA, whereas Class 2 genes are gradually activated from early neurula and onward but not significantly before (Figure 5C, Figure S5A). These data suggest that Hdac1 regulates both transcriptionally active and silent genes at gastrulation.

**Figure 5.**
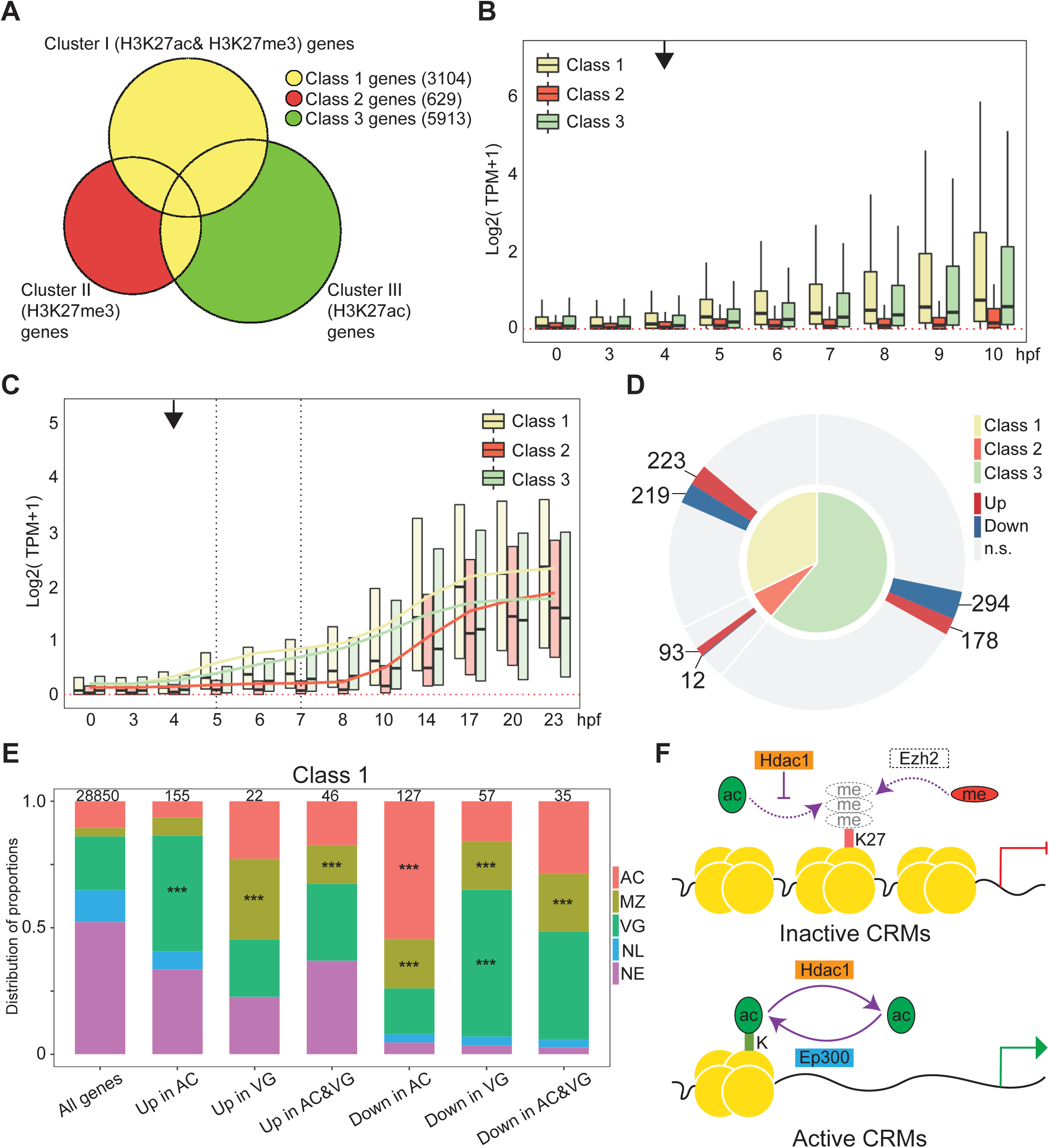
Hdac1 regulates germ layer transcriptomes both in time and space. (**A**) Venn diagram comparing genes associated with Hdac1 CRM clusters (Clusters I, II, and III in Figure 3C). Class 1 are genes closest to Cluster I peaks plus overlapped genes in both Class 2 and 3; Class 2 are unique genes closest to Cluster II peaks; Class 3 are unique genes closest to Cluster III peaks. (**B**) Time-course TPM expression of zygotic genes in Classes 1, 2, and 3 up to 10 hours post-fertilization (hpf, late st12.5). Red dotted line denotes zero. Black arrow denotes the onset of ZGA. (**C**) Time-course TPM expression of zygotic genes in Classes 1, 2, and 3 up to 23 hpf (tailbud st26). Black vertical dotted lines denote the time window when Hdac1 binding is examined. Trend lines for each class are generated by connecting mean TPM values at each time point. (**D**) Nested pie chart illustrating the proportions of genes affected by TSA in each class. n.s.: not significant. (**E**) Spatial expression pattern of Class 1 genes affected by TSA in proportions. The total number of genes in each category is listed at the top of each bar. Only *** denoting *p*< 0.001 (Fisher’s exact test) is shown. AC: animal cap, presumptive ectoderm; MZ: marginal zone, presumptive mesoderm; VG: vegetal mass, presumptive endoderm; NL: non-localized genes; NE: non-expressed genes. Parameters of spatial categories are detailed in Supplemental Methods. (**F**) Model of Hdac1 functioning at both inactive and active CRMs: on inactive CRMs, H3K27 residue is maintained as unmodified by Hdac1, which may be subjected to H3K27me3-mediated suppression; on active CRMs, the state of histone acetylation is dynamically modulated by Ep300-and-Hdac1-mediated acetylation-deacetylation cycles.

To understand how HDAC activity affects the expression of nearby genes, we performed RNA sequencing (RNA-seq) using early gastrula AC or VG explants treated with or without TSA (Figure S4B). Differential gene expression analyses revealed a greater number of differentially regulated genes in AC (1401 genes) than in VG (471 genes) suggesting that the ectodermal lineage is more susceptible to the loss of HDAC activity (Figure S5B). Many genes known to function during the germ layer development are affected when HDAC activity is inhibited. Genes affected by TSA treatment are affected similarly by VPA treatment indicating that identified genes are not due to non-specific effects of TSA (Figure S5C). We found that HDACs have many overlapping and non-overlapping gene targets between AC and VG suggesting that HDACs confer both specific and ubiquitous activities in different germ layers (Figure S5D). Gene ontology analyses revealed that genes affected by HDAC inhibition primarily function in early embryonic development such as pattern specification, cell fate commitment, regionalization, tissue morphogenesis and many other processes. (Figure S5E, S5F).

### Integrity of germ-layer specific transcriptomes is maintained by Hdac1 activity temporally and spatially

We attempt to correlate how the expression of different classes of Hdac1 bound genes (Classes 1-3 in Figure 5A) is affected by HDAC inhibition. Differentially regulated genes in Class 2 (whose CRMs are only marked by H3K27me3) are primarily (>85%) activated when HDAC activity is inhibited (Figure 5D). Given that Class 2 genes are not transcribed until after gastrulation normally (Figure 5C), premature activation of these genes at early gastrula upon HDAC inhibition supports the idea that Hdac1 suppresses the transcription of these genes temporally. This is consistent with observed histone hyperacetylation at surrounding repressive CRMs upon HDAC inhibition (Figure 4E). Furthermore, we observed that differentially regulated genes in Class 1 (whose CRMs are marked by a mixture of H3K27ac and/or H3K27me3) and Class 3 (whose CRMs are only marked by H3K27ac) can either be up- or down-regulated upon HDAC inhibition (Figure 5D). We speculate that HDAC activity is spatially employed by genes in Class 1 and 3.

To gain insights into the mechanism of spatial gene regulation by Hdac1, we analyzed the spatial expression patterns of all genes at early gastrula (Blitz et al., 2017) and subdivided genes into 5 categories (AC, MZ, and VG: genes enriched in ectoderm, mesoderm, and endoderm lineages, respectively. NL: uniformly expressed genes, and NE: lowly or not expressed genes). We then examined the endogenous spatial expression patterns of Class 1 genes affected by HDAC inhibition. Compared to spatial transcriptomic profiles of wide-type embryos, genes normally expressed in endoderm are significantly up-regulated in AC, and genes normally expressed in mesoderm are significantly up-regulated in VG upon HDAC inhibition (Figure 5E). This finding reveals that Hdac1 prevents misactivation of genes in respective germ layers. We highlight that misactivated genes are highly correlated with histone hyperacetylation of surrounding CRMs. For example, VG CRMs are hyperacetylated in AC upon HDAC inhibition (VG CRMs in Figure 4F), which in turn leads to the misactivation of endodermal genes in AC. Surprisingly, we found that genes normally expressed in ectoderm and mesoderm are significantly down-regulated in AC, and genes normally expressed in mesoderm and endoderm are significantly down-regulated in VG upon HDAC inhibition (Figure 5E). Contrary to the repressive role of Hdac1, this finding suggests that Hdac1 positively influences the transcription of active genes in each germ layer. We propose that the state of histone acetylation on CRMs associated with active genes is dynamic, and disruption to such equilibrium impairs normal transcription activities. Based on the quantitative analysis of pan-H3Kac profiles upon TSA treatment, we found that Hdac1 bound CRMs gain increased levels of pan-H3Kac globally including on already acetylated CRMs (Figure S4G, 4F). This TSA-induced excessive histone acetylation on active CRMs may contribute to the attenuated expression of associated active genes in respective germ layers. A similar trend is observed among TSA-responsive Class 3 genes (Figure S5G). These data demonstrate a spatial regulation where Hdac1 not only prevents aberrant activation of silent genes, but also sustains proper gene expression levels in each germ layer.

Since both Class 1 (whose CRMs are marked by a mixture of H3K27ac and/or H3K27me3) and Class 3 (whose CRMs are only marked by H3K27ac) genes show germ-layer specific pan-H3Kac signals and transcriptomic profiles, we sought to examine the differences between these two classes. Class 1 genes display a higher variability in expression levels between different germ layers compared to Class 3 genes (Figure S5H). This indicates that the expression of genes undergoing active H3K27me3 suppression is more spatially restricted; thus, their functions may be more intimately associated with germ layer determination. Altogether, these results show that Hdac1 maintains the integrity of germ layer genes both in time and space during gastrulation.

## Discussion

Here we defined a critical role for Hdac1 during early *Xenopus* embryogenesis. Our findings demonstrate a close link between homeostasis of histone acetylation and transcriptional activities in developing embryos. Progressive binding of Hdac1 to the genome shapes the zygotic histone acetylome, thereby reinforcing a proper germ-layer specific transcriptome, both in time and space. Thus, Hdac1 is an essential epigenetic regulator in the control of embryonic cell identity and lineage. We propose that TFs inducing differentiation programs exploit the activity of Hdac1 to confine the expression of zygotic genes.

### Gradual binding of Hdac1 coincides with ZGA

The genome-wide binding of Hdac1 begins at mid blastula and gradually accumulates at thousands of loci. Such progressive binding of Hdac1 around ZGA raises the question of whether Hdac1 recruitment requires zygotic factors during these stages. Surprisingly, ChIP-seq analyses of Hdac1 from α-amanitin-injected embryos show that zygotic transcription is dispensable for Hdac1 recruitment to the genome, at the very least, up to late blastula (Figure 2A, S2B). Recent work in yeast shows that active transcription is required to shape histone acetylation patterns largely due to a direct role of RNAPII in recruitment and activation of H4 HATs but not HDACs (Martin et al., 2021). Our result agrees with the notion that Hdac1 recruitment is not directed by on-going transcription. We, therefore, speculate that maternal factors instruct early Hdac1 recruitment. Our results reveal a positive binding correlation between Hdac1 and maternal Foxh1/Sox3, inferring the importance of these two maternal TFs in Hdac1 early recruitment. Though maternal TFs such as Foxh1, Vegt, and Otx1 are shown to bind the genome as early as the 32∼64-cell stage (Charney et al., 2017; Paraiso et al., 2019), the genome-wide binding of Hdac1 begins at blastula and is not significantly widespread until early gastrula (Figure 1C). It is also known that Hdac1 containing complexes bind to pre-existing epigenetic marks such as DNA methylation and H3K4me3 (Wade et al., 1999; Lee et al., 2018). A previous study in *Xenopus* demonstrates that H3K4me3 is primarily instructed maternally until late gastrula (Hontelez et al., 2015). Therefore, maternally instructed histone modifications may also play a role in Hdac1 recruitment.

### Hdac1 functions differently on active versus inactive CRMs

One perplexing finding is that Hdac1 occupies both active and repressive genomic loci (Figure 3A, 3B). Hdac1 binds to repressive genomic regions that are facultative but not constitutive heterochromatin. Though histone modifications underlying constitutive heterochromatin have been shown to regulate developmental genes (Riddle et al., 2011; Wang et al., 2018; Methot et al., 2021), the profile of H3K27me3 is largely different (∼90% non-overlapping peaks) from profiles of H3K9me2, 3, and H4K20me3 in early *Xenopus* development (data not shown, partially shown in van Kruijsbergen et al., 2017). These observations suggest that Hdac1-mediated suppression is largely dictated by developmental programs. In contrary to commonly accepted repressor function of Hdac1, the binding of Hdac1 at active genomic regions is surprising. However, our findings are consistent with previous studies in yeast and mammalian cell culture suggesting a dynamic equilibrium of histone acetylation at active loci (Kurdistani et al., 2002; Wang et al., 2002; Wang et al., 2009; Kidder et al., 2012). Hence, we hypothesize that Hdac1 has different functions on active versus inactive CRMs in early embryos.

To test the *in vivo* function of HDACs, we blocked their endogenous activity using an inhibitor and quantitatively examined the changes of general H3 acetylation (pan-H3Kac) upon HDAC inhibition. First, an increase of pan-H3Kac is observed across Hdac1-bound CRMs, including active CRMs, consistent with the canonical enzymatic activity of Hdac1 (Figure S4G). Second, repressive CRMs marked by H3K27me3 undergo drastic H3 hyperacetylation compared to active CRMs marked by H3K27ac suggesting a HDAC-activity dependent suppression of repressive CRMs (Figure 4E). Third, germ-layer specific pan-H3Kac profiles are disrupted indicating the importance of Hdac1 in defining spatial patterns of histone acetylation (Figure 4F, S4J). Based on these results, we propose a dual function model for Hdac1. On the one hand, Hdac1 prevents histone acetylation at inactive CRMs, thereby preserving H3K27 as unacetylated (Figure 5F, inactive CRMs model). Interestingly, H3K27me3 is not always imposed on inactive CRMs. For instance, active CRMs (Cluster III, only H3K27ac) are spatially modified with pan-H3Kac but are not subjected to H3K27me3 (Figure 3D, 4D, S4I, S4J). This suggests that HDAC-mediated histone deacetylation and Polycomb-mediated histone methylation are not coupled at inactive CRMs. On the other hand, Hdac1 participates in dynamic histone acetylation-deacetylation cycles at active CRMs (Figure 5F, active CRMs model). Although we did not directly test the co-binding of HATs and HDACs, CRMs may be simultaneously bound since (1) the binding profiles of Ep300 and Hdac1 mostly overlap (Figure S3A), and (2) pan-H3Kac signals increase at all Hdac1 peaks including active CRMs, upon HDAC inhibition (Figure S4G). Presumably, active CRMs are maintained in a state of dynamic equilibrium. This model is in accordance with a previous study demonstrating that HATs and HDACs simultaneously participate in histone acetylation cycles, which initiate and reset chromatin between rounds of transcription (Wang et al., 2009).

### Hdac1 safeguards misactivation of genes both in time and space

We attempted to correlate the activity of CRMs with the transcriptional activity of potential target genes. Genes associated with repressive CRMs (H3K27me3 only) are mainly inactive until neurula (Figure 5B, 5C). More than 85% of these genes are prematurely activated upon HDAC inhibition suggesting that Hdac1 maintains the state of histone hypoacetylation on repressive CRMs, thereby preventing premature expression of genes. Moreover, genes associated with active (H3K27ac only) and heterogeneous (both H3K27ac and H3K27me3) CRMs are misactivated in different germ layers when HDAC activity is blocked (Figure 5E, S5G). This indicates that Hdac1 safeguards differential histone acetylation states in each germ layer (Figure 4F, S4J), resulting in a proper spatial transcriptome. We did not directly address whether hypoacetylated heterogeneous CRMs are subjected to H3K27me3. However, a previous study showed that H3K27me3 is spatially deposited in line with spatial patterns of gene expression at late gastrula (Akkers et al., 2009). We predict that heterogenous CRMs are differentially marked by opposing H3K27me3 or acetylation in different germ layers. In summary, Hdac1 preserves the histone hypoacetylation state of inactive CRMs resulting in gene suppression both in time and space, which follows the canonical transcriptional corepressor role of Hdac1.

### Cyclical histone acetylation sustains germ layer gene transcription

Our study reveals an unexpected role for Hdac1 in sustaining active gene expression during germ layer formation. Within both ectoderm and endoderm, the expression of TSA down-regulated genes associated with either active (H3K27ac only) or heterogeneous (H3K27ac and H3K27me3) CRMs are enriched in their respective germ layers (Figure 5E, S5G). This suggests a paradoxical activator role for Hdac1, which is also reported in previous studies (Vidal and Gaber, 1991; Zupkovitz et al., 2006; Baltus et al., 2009; Hughes et al., 2014; Rao and LaBonne, 2018). We speculate that utilization of HDAC activities at active genomic loci is a general mechanism as seen in examples of *Xenopus* ectoderm and endoderm lineages which deploy distinct gene regulatory networks. Based on our findings, we propose that a dynamic equilibrium between acetylation and deacetylation is essential to sustain gene transcription. The function of HDACs on active genomic regions has been elucidated in several contexts. In yeast, cotranscriptional methylation (H3K36me3, H3K4me2) recruits HDAC containing complexes (Rpd3S, Set3C) to suppress intragenic transcription and delay induction of genes that overlap non-coding RNAs (Carrozza et al., 2005; Keogh et al., 2005; Li et al., 2007; Kim and Buratowski, 2009; Kim et al., 2012; Heo at al., 2021). Genetic deletion of Set3C affects transcript levels only in altered growth conditions (Lenstra et al., 2011), consistent with the notion that cyclical histone acetylation act as a mechanism to regulate dynamics of RNA induction and fidelity of transcription. Studies in metazoans show a conserved mechanism. HDAC1 can be targeted by p300 to transcribing genes through a direct interaction (Simone et al., 2004). Simultaneous binding of both HATs and HDACs at active genomic regions is shown in T cells (Wang et al., 2009). Inhibition of both DNA methyltransferases and HDACs induces cryptic transcription in lung cancer cells (Brocks et al., 2017). Down-regulated genes upon HDAC inhibition exhibit high levels of cryptic transcripts during mouse cardiogenesis (Milstone et al., 2020). These findings suggest a role for HDAC activity in transcriptional fidelity.

Why does excessive histone acetylation due to disruption of cyclical histone acetylation lead to reduced transcription instead of elevated transcription? We observed that active (H3K27ac only) and heterogeneous (both H3K27ac and H3K27me3) CRMs are excessively acetylated (Figure 4E, S4G) following HDAC inhibition, which results in the reduced expression of CRM-associated genes within their respective germ layers (Figure 5E, S5G). This leads us to speculate that excessive histone acetylation interferes the activity of the transcriptional machinery. A recent study shows that excessive histone acetylation on chromatin induced by inhibition of HDAC1, 2, and 3 leads to increased aberrant contacts and reduced native contacts between super-enhancer loops (Gryder et al., 2019). This suggests that the excessive histone acetylation impairs active transcription by altering chromatin interactions. Alternatively, excessive histone acetylation can alter the binding of acetyl-histone readers. H4 polyacetylation induced by HDAC inhibition is shown to be preferentially bound by BRD proteins (such as BRD4), thereby sequestering these factors away from active genes (Slaughter et al., 2021). Therefore, HDACs safeguards the function of normal acetyl-histone readers. Further investigation of cyclical histone acetylation regulating developmental programs is needed in the context of germ layer specification.

## Materials and Methods

### Animal Model and Subject Details

*Xenopus tropicalis* embryos were obtained by *in vitro* fertilization according to Ogino et al. (2006) and staged according to Nieuwkoop and Faber (1994). All embryos were cultured in 1/9X Marc’s modified Ringers (MMR) at 25 °C. For HDAC inhibition, 4-cell stage embryos were immersed in 1/9X MMR containing 1). 100nM TSA (Esmaeili et al., 2020) or DMSO; or 2). 10mM VPA (Rao and LaBonne. 2018) or H_2_O. For α-amanitin injection, each 1-cell stage embryo was injected with 6pg of α-amanitin (Hontelez et al., 2015). For embryo dissection, embryos were dissected at the late blastula stage (6 hpf), and explants were cultured to the early gastrula (7 hpf). Animals were raised and maintained following the University of California, Irvine Institutional Animal Care Use Committee (IACUC). Animals used were raised in the laboratory and/or purchased from the National *Xenopus* Resource (RRID: SCR_013731).

### Western Blotting

Embryos were homogenized in 1X RIPA (50mM Tris-HCl pH7.6, 1% NP40, 0.25% Na-Deoxycholate, 150mM NaCl, 1mM EDTA, 0.1% SDS, 0.5mM DTT) with protease inhibitors (Roche cOmplete) and centrifuged twice at 14,000 rpm. The supernatant was then subjected to western blotting using anti-HDAC1 (Cell Signaling, 34589S), anti-HDAC2 (Genetex, GTX109642), and anti-Tubulin (Sigma, T5168). For histone modifications, acid-extracted histone lysate was prepared accordingly (Shechter et al., 2007) and subjected to western blotting using anti-H3K9ac (Cell Signaling, 9649), H3K14ac (Cell Signaling, 7627), H3K18ac (Cell Signaling, 13998), H3K27ac (Cell Signaling, 8173), H3K56ac (Cell Signaling, 4243), and H4K20ac (Active Motif, 61531).

### Chromatin Immunoprecipitation (ChIP)

ChIP protocol was performed as described (Chiu et al., 2014). Antibodies used for ChIP were anti-HDAC1 (Cell Signaling, 34589S, 1:100), anti-H3K18ac (Cell Signaling, 13998, 1:100), and anti-Sox3 (Zhang et al., 2003). ChIP-seq libraries were constructed using the NEBNext Ultra II DNA Kit (NEB, E7645).

For sequential ChIP, the first round of ChIP was performed as described and eluted in 1X TE containing 1% SDS at 37 °C for 30 minutes. The eluate was diluted with 1X RIPA (without SDS) 10 times and subjected to the second round of ChIP as described (Desvoyes et al., 2018). Real-time quantitative PCR (RT-qPCR) was performed using Power SYBR Green PCR master mix (Roche) to quantify the DNA recovery compared to ChIP input DNA at one embryo equivalency (percent input). The error among technical replicates was calculated using the rule of error propagation. Primer sequence information is provided in Table S1.

For dissected pan-H3Kac ChIP, spike-in chromatin (Active Motif, 53083) was added to the chromatin of dissected tissues at a ratio of 1:35. Mixed chromatin was then subjected to ChIP with 5ug anti-panH3Kac (Active Motif, 39139) and 1ug of anti-H2Av (Active Motif, 61686) and followed as described.

All experiments were performed in two independent biological replicates unless noted. Sequencing was performed using the Illumina NovaSeq 6000 and 100bp single-end reads or 100bp paired-end reads were obtained. Computational analysis of ChIP-seq is described in the Supplemental Methods.

### RNA-seq and Gene Expression Assays

Total RNA from dissected tissues was extracted using Trizol as described (Amin et al., 2014). mRNA was then isolated using NEBNext PolyA mRNA Magnetic Isolation Module (NEB E7490S). Sequencing libraries were prepared using NEBNext Ultra II RNA library prep kit (NEB E7770S) and sequenced by the Illumina NovaSeq 6000 with 100bp paired end reads. All experiments were performed in two independent biological replicates. Computational analysis of RNA-seq is described in the Supplemental Methods.

For RT-qPCR, reverse-transcription assay was performed using Maxima reverse transcriptase (Thermo Fisher EP0741). RT-qPCR was performed using LightCycler 480 SYBR Green I master mix (Roche). For quantification of gene expression, the 2^ -ΔΔCt method was used. The expression level of *drosha* or *rsrc1* was used for normalization. The error among technical replicates was calculated using the rule of error propagation. Primer sequence information is provided in Table S3.

### Data Accessibility

Raw and processed RNA-seq and ChIP-seq datasets generated from this study are available at NCBI Gene Expression Omnibus using the accession GSE198378. Publicly available datasets used in this study are available at NCBI Gene Expression Omnibus using the accession GSE56000 (Gupta et al., 2014; H3K27ac ChIP-seq), GSE67974 (Hontelez et al., 2015; Ep300, H3K9ac, H3K4me1, H3K4me3, H3K36me3, H3K9me2, H3K9me3, H3K27me3, H4K20me3 ChIP-seq and st12 RNA Pol2 ChIP-seq), GSE65785 (Owens et al., 2016; temporal profiling of RNA-seq), GSE85273 (Charney et al., 2017; st9 Foxh1 ChIP-seq, st7, 8, 9& 10.5 RNA Pol2 ChIP-seq), GSE81458 (Blitz et al., 2017; st10.5 dissected germ-layer RNA-seq).

## Competing interest statement

The authors declare no competing interests.

## Acknowledgments

We thank Drs. Kyoko Yokomori (University of California, Irvine), Yongsheng Shi (University of California, Irvine), and current Cho lab members for insightful comments on this study. We thank the Genomic High Throughput Facility at the University of California, Irvine for sequencing services. We also thank the Research Cyberinfrastructure Center at the University of California, Irvine for the ongoing support of High Performance Community Computing Clusters. This work is supported by NIH R01GM126395, R35GM139617 and NSF 1755214 to K.W.Y.C..

## Author contributions

Conceptualization & Methodology: J.J.Z., K.W.Y.C.; Investigation: J.J.Z., J.S.C., H.H., W.W.; Resources: W.W., I.L.B., K.W.Y.C.; Software, Formal analysis, Data curation & Visualization: J.J.Z.; Writing-original draft: J.J.Z., I.L.B., K.W.Y.C.; Supervision & Funding acquisition: K.W.Y.C.

## Ethics

All animals were manipulated according to an approved institutional animal care and use committee (IACUC) protocols (AUP-21-068) of the University of California, Irvine. Every effort (such as Tricaine anesthesia) was made to minimize suffering of animals. All animal husbandry is guided by methods developed by the National Xenopus Resource (Marine Biological Laboratory, Woods Hole, MA).

## Supplementals

### Supplemental Methods

#### ChIP-seq Analysis

All sequencing data were aligned to *Xenopus tropicalis* v10.0 genome (http://www.xenbase.org/, RRID:SCR_003280) using Bowtie2 v2.4.4 (Langmead and Salzberg, 2012). PCR duplicates were removed using Samtools v1.10 (Li et al., 2009). ChIP-seq signals were visualized using IGV v2.11.3 (Robinson et al., 2011) after concatenating two biological replicates when available. Irreproducibility discovery rate (IDR) analysis (Li et al., 2011) was used to identify high-confidence peaks called by Macs2 v2.7.1 (Zhang et al., 2008) against the stage-matched input (Charney et al., 2017) between two biological replicates according to ENCODE3 ChIP-seq pipelines (IDR threshold of 0.05) (https://docs.google.com/document/d/1lG_Rd7fnYgRpSIqrIfuVlAz2dW1VaSQThzk836Db99c/edit).

For dissected pan-H3Kac ChIP, **all second replicates** were downsampled to 25% by random sampling to a comparable sequencing depth for later analysis. *Drosophila* H2Av peaks are generated from overlapped peaks with two published S2 cell samples (Weber et al., 2014; Tettey et al., 2019). Normalization factors were then calculated based on reads that mapped to *Drosophila* H2Av peaks for each ChIP-seq sample as used in Egan et al., 2016. When replicates or dissected samples are merged for analyses, normalization factors are calculated as the sum of Ratio_frog_ *Ratio_fly_. Detailed normalization factors used are listed in Table S2.

#### RNA-seq Analysis

All sequencing samples were aligned using STAR v2.7.3a (Dobin et al., 2013) to *Xenopus tropicalis* genome v10.0 (http://www.xenbase.org/, RRID:SCR_003280) to obtain raw read counts. RSEM v1.3.3 (Li and Dewey, 2011) was used to calculate expression values in transcripts per million (TPM). Differentially expressed genes were identified using edgeR v3.36.0 (Robinson et al., 2010) with parameters, greater than 2-fold change and less than 0.05 false discovery rate (FDR, also known as the adjusted p-value), in R v4.1.2 (R Core Team, 2021). Metascape (Zhou et al., 2019) was used to perform gene ontology enrichment analyses with default parameters (min overlap= 3, p-value cutoff= 0.01, and min enrichment= 1.5).

#### Additional Bioinformatics and Statistical Analyses

Samtools v1.10 (Li et al., 2009) was used to convert between SAM and BAM files. DeepTools v3.5.0 (Ramírez et al., 2014) was used to generate: (1) ChIP-seq signal track (bigwig files) normalized by reads per genomic content (-RPGC) at the bin size of 1bp; (2) heatmaps around peak summits normalized by Bins Per Million mapped reads (-BPM) at the bin size of 50bps; (3) signal profile along the gene bodies normalized by -BPM at the bin size of 50bps; (4) Pearson correlation between ChIP-seq samples at peaks. Homer v4.10 (Heinz et al., 2010) was used to annotate genomic features of ChIP peaks. Bedtools v2.29.2 (Quinlan and Hall, 2010) was used to determine peak overlaps among ChIP-seq peaks and obtain counts of reads at ChIP-seq peaks. CentriMo (Bailey and Machanick, 2012) was used to perform local motif enrichment analysis. Welch Two Sample t-test in R v4.1.2 was used to determine the statistical significance between groups. Cohen’s d (effect size) was calculated using lsr v0.5.2 package in R v4.1.2 (R Core Team, 2021). Time-course gene expression was obtained from ribosomal RNA-depleted RNA-seq data (Owens et al., 2016). The expression in TPM was calculated as outlined above. Spatial gene expression at early gastrula was obtained from RNA-seq of 5 dissected tissues (Blitz et al., 2017) consisting of the animal cap (ectoderm), the dorsal marginal zone (dorsal mesoderm), the lateral marginal zone (lateral mesoderm), the ventral marginal zone (ventral mesoderm), and the vegetal mass (endoderm). The expression in TPM was obtained as outlined above. Fisher’s Exact test (alternative = “greater”) in R v4.1.2 (R Core Team, 2021) was used to determine the significance of the proportional enrichment between groups.

#### Categorical Analyses

Spatial categorization of CRMs is defined as below: AC CRMs represent CRMs whose pan-H3Kac signals in AC is 2-fold higher than in VG; VG CRMs represent CRMs whose pan-H3Kac signals in VG is 2-fold higher than in AC; ubiquitous CRMs represents the remaining CRMs whose pan-H3Kac signals do not exceed 2-fold enrichment in either of the two examined germ layers. For temporal gene expression analysis, (strictly) zygotic genes are determined by removing genes whose expression levels are greater than 1 TPM during the first 2 hours post-fertilization. Spatial categorization of genes at early gastrula stage: the average TPM between 3 dissected mesoderm tissues (dorsal, marginal, and lateral marginal zones) was used to represent the expression of mesoderm. Genes with the expression in any dissected tissue less than 1 TPM were considered not expressed. Genes with the coefficient of variance of TPM less than 0.1 (10%) were considered evenly expressed. The remaining genes with localized expression were assigned to a germ layer based on the maximum TPM.

---

**Figure S1.**
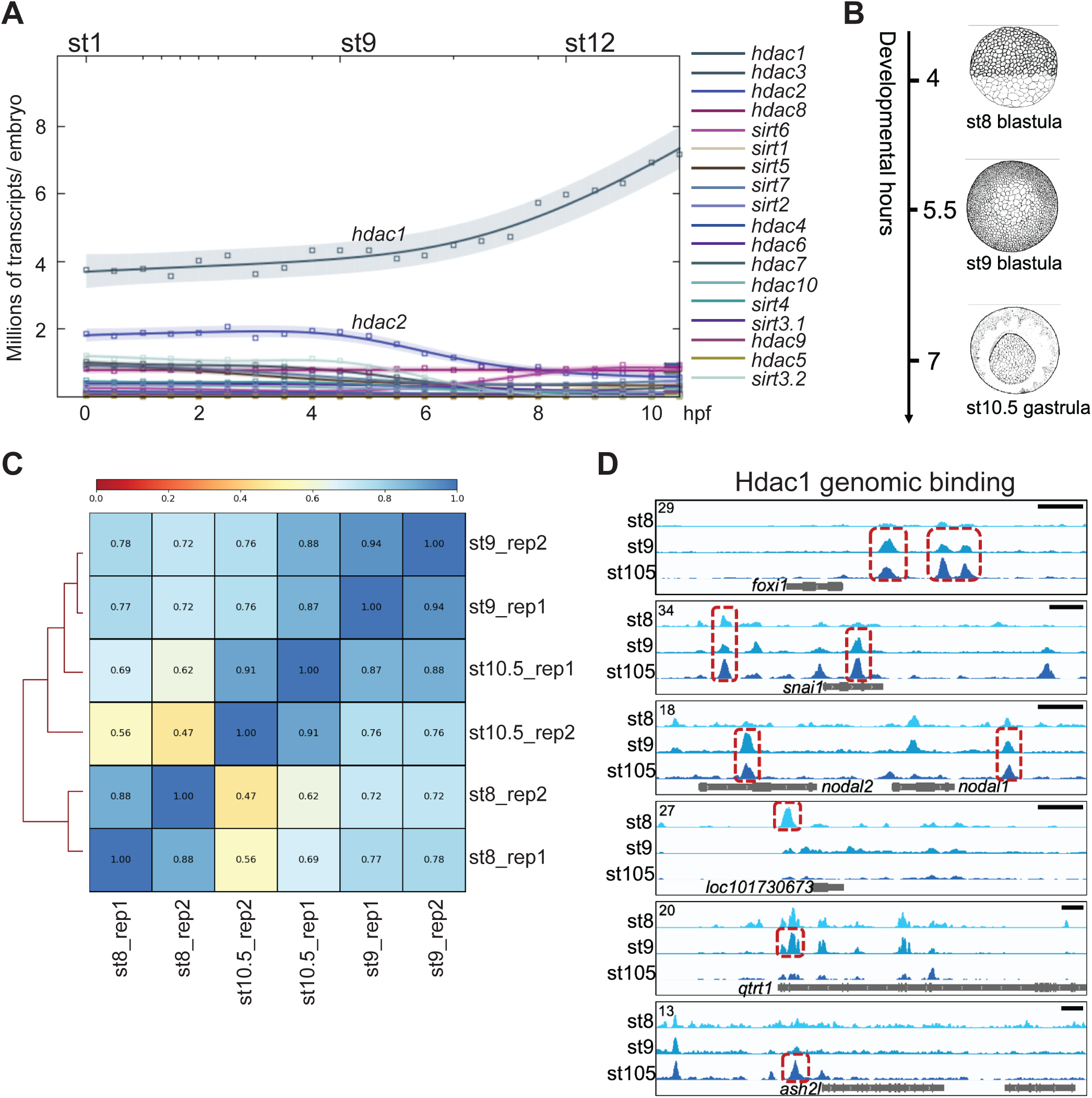
Hdac1 genomic occupancy in early *Xenopus* development. (**A**) Time-course expression levels of HDACs revealed by ribo-depleted RNA-seq (rdRNA-seq) in *Xenopus tropicalis*. (**B**) Graphic illustration of examined stages during early *Xenopus* development: st8 mid blastula, st9 late blastula, and st10.5 early gastrula (Nieuwkoop and Faber, 1994). (**C**) Pearson correlation analyses of intra- and inter-stage Hdac1 ChIP-seq samples. (**D**) Genome browser visualization of Hdac1peaks on *foxi1*, *snai1*, *nodal1*, *nodal2*, *loc101730673*, *qtrt1*, and *ash2l*. Red boxes represent IDR peaks passing IDR threshold (see Materials and Methods) on listed genes (not all IDR peaks are boxed). Y-axis values represent scaled track height. Black bars denote an interval of 2kb.

**Figure S2.**
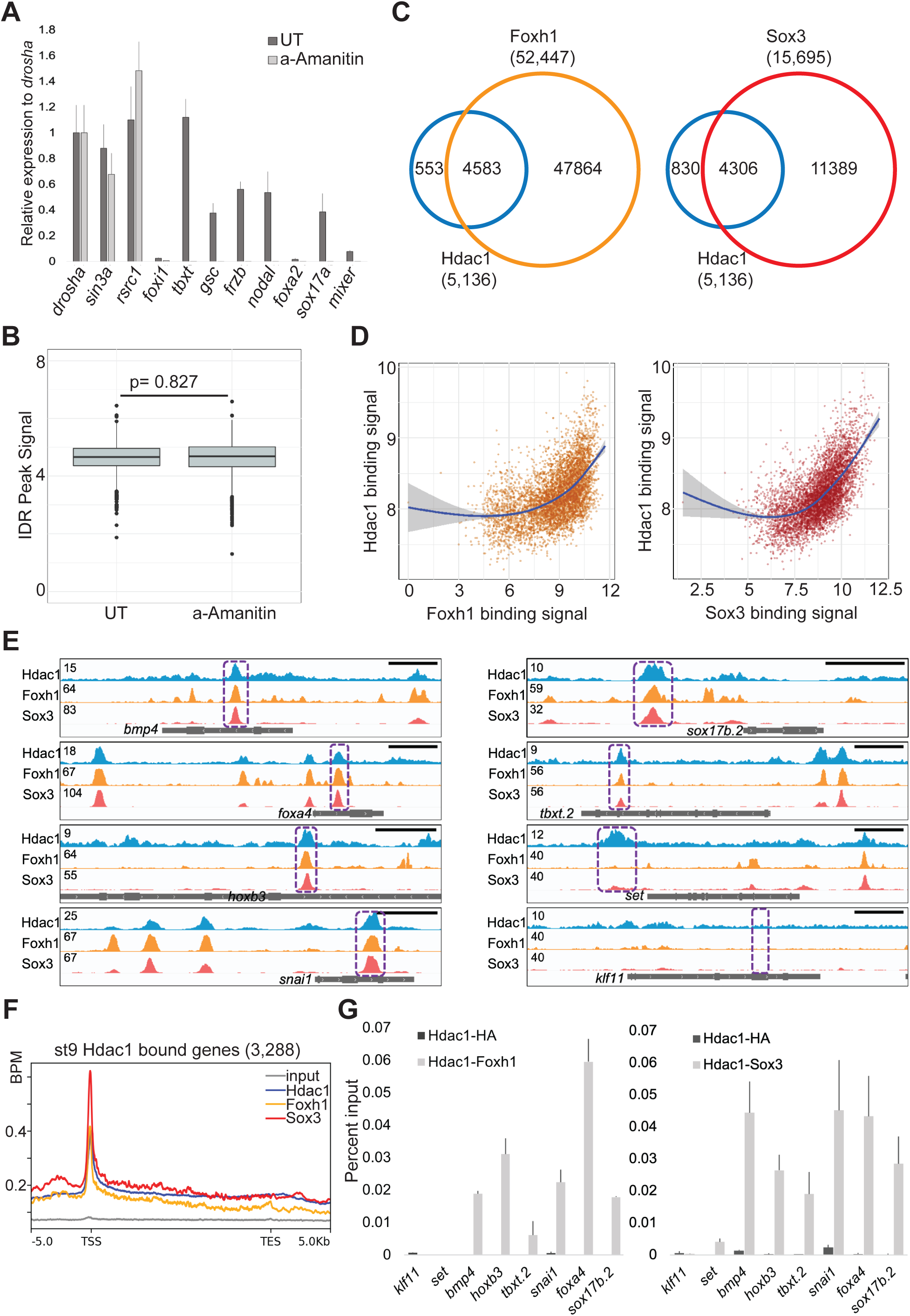
Hdac1 binding to genome is instructed maternally. (**A**) RT-qPCR analysis of maternal (maternal-zygotic) genes (*drosha*, *sin3a*, and *rsrc1*) and zygotic genes (*foxi1*, *tbxt*, *gsc*, *frzb*, *nodal*, *foxa2*, *sox17a*, and *mixer*) showing a high efficacy of α-amanitin blocking zygotic transcription. The error bars represent the variation from 2 technical replicates. (**B**) Box plot illustrating Hdac1 signal enrichment at st9 Hdac1 IDR peaks between uninjected and α-amanitin injected embryos. Y-axis values represent log2(reads to IDR peaks/ (peak size* read depth at IDR peaks) +1). *p*-Value is calculated by two-tailed Student’s *t*-test. (**C**) Venn diagram showing the overlap between st9 IDR peaks of Foxh1 and Hdac1; Sox3 and Hdac1. (**D**) Scatter plots representing a positive correlation between binding signals of Foxh1 and Hdac1; Sox3 and Hdac1. The binding signal is calculated as log2(reads mapped to st9 Hdac1 IDR peaks per kb +1). Blue lines depict linear regression curves generated from a generalized additive model. Grey areas denote a 95% confidence interval. (**E**) Genome browser visualization of Hdac1, Foxh1, and Sox3 signals on *bmp4*, *foxa4*, *hoxb3*, *snai1*, *sox17b.2*, *tbxt.2*, *set*, and *klf11* genes. Purple boxes represent genomic regions examined in sequential ChIP-qPCR experiments. Y-axis values represent scaled track height. Black bars denote an interval of 2kb. (**F**) Distributions of Hdac1, Foxh1, and Sox3 ChIP-seq signals within the intervals of 5 kb upstream of gene model 5’ ends, gene bodies (normalized for length), and 5 kb downstream of gene model 3’ ends. The signal of st9 input DNA ChIP-seq is used as a negative control. Y-axis values represent reads quantified by bins per million (BPM) at a bin size of 50 bp. (**G**) Reciprocal sequential ChIP-qPCR analyses to Figure 2D& 2E assessing Foxh1 and Hdac1 co-bound regions; Sox3 and Hdac1 co-bound regions. anti-HA is used as a negative control. The error bars represent the variation from 2 technical replicates.

**Figure S3.**
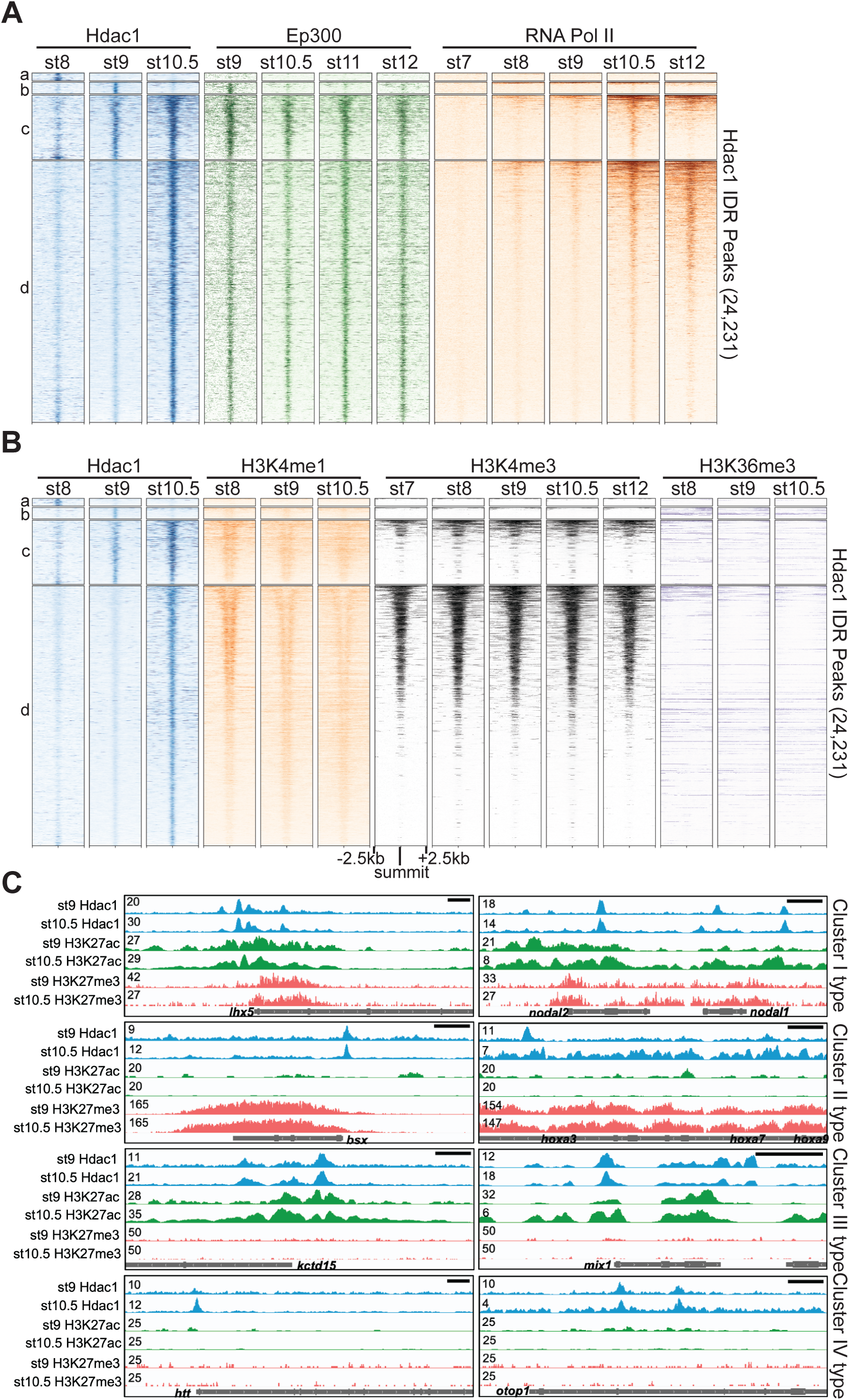
Hdac1 bound CRMs are marked with distinct epigenetic signatures. (**A, B**) Clustered heatmaps showing signals across several stages of (**A**) Ep300, RNA polymerase II, and (**B**) various active histone methylation marks on Hdac1 peaks. The signals are shown in a window of 5kb centered on the summits of Hdac1 peaks. The signal density within each cluster is ordered by RNA polymerase II in (**A**) and by all samples in (**B**) by descending order. Each cluster corresponds to the same region in Figure 1C. (**C**) Genome browser visualization of Hdac1, H3K27ac, and H3K27me3 signals on genes representing Hdac1 CRM clusters: Cluster I (lhx*5*, *nodal1*, *nodal2*), Cluster II (*kctd15*, *mix1*), Cluster III (*bsx*, *hoxa3*, *hoxa7*, *hoxa9*) and Cluster IV (*htt*, *otop1)*. Y-axis values represent scaled track height. Black bars denote an interval of 2kb.

**Figure S4.**
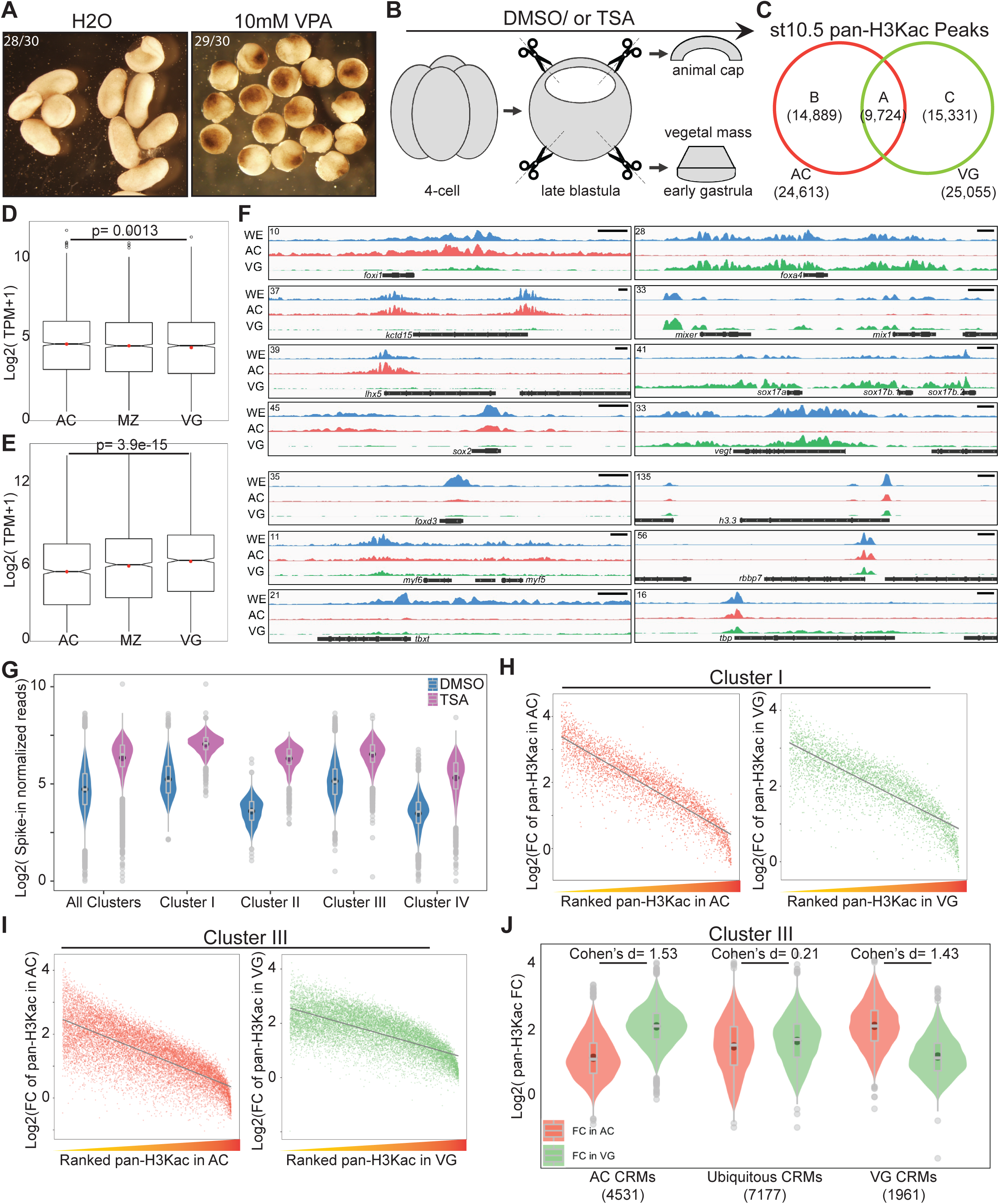
Germ-layer specific H3 acetylomes require HDAC activity. (**A**) Embryos treated with 10mM valproic acid (VPA) display gastrulation defects 24 hours post-fertilization. (**B**) A schematic diagram of dissection experiments to separate animal cap tissues (AC) and vegetal mass tissues (VG) for RNA-seq or ChIP-seq. Late blastula stage explants were cultured ∼1-1.5hrs to the early gastrula stage before being harvested for analyses. (**C**) Venn diagram comparing pan-H3Kac peaks in AC and VG. (**D, E**) Transcripts per million (TPM) expression of genes associated with (**D**) AC specific pan-H3Kac and (**E**) VG specific pan-H3Kac peaks across three germ layers. *p*-values are calculated by Student’s *t*-test. (**F**) Genome browser visualization of pan-H3Kac signals at known ectodermal (*foxi1*, *kctd15*, *lhx5*, *sox2*), mesodermal (*foxd3*, *myf5*, *myf6*, *tbxt*), endodermal (*foxa4*, *mixer*, *mix1*, *sox17a*, *sox17b.1*, *sox17b.2*, *vegt*) and “housekeeping” (*h3.3*, *rbbp7*, *tbp*) genes in whole embryos, AC, and VG. Y-axis values represent scaled track height. Black bars denote an interval of 2kb. (**G**) Spike-in normalized pan-H3Kac signals across Hdac1 CRM clusters (clusters in Figure 3C) in DMSO- or TSA-treated samples. Signals of pan-H3Kac are combined from experiments done in AC and VG. (**H, I**) Scatter plots representing a negative correlation between fold changes of pan-H3Kac signals and ranked endogenous pan-H3Kac signals in AC or VG at both (**H**) Cluster I and (**I**) Cluster III Hdac1 CRM clusters (clusters in Figure 3C). Grey lines depict linear regression curves generated from a linear model. Both x- and y-axes’ values are log2 transformed. (**J**) Fold changes (FC) of pan-H3Kac signals at Cluster III of Hdac1 CRM clusters categorized spatially (described in Methods).

**Figure S5.**
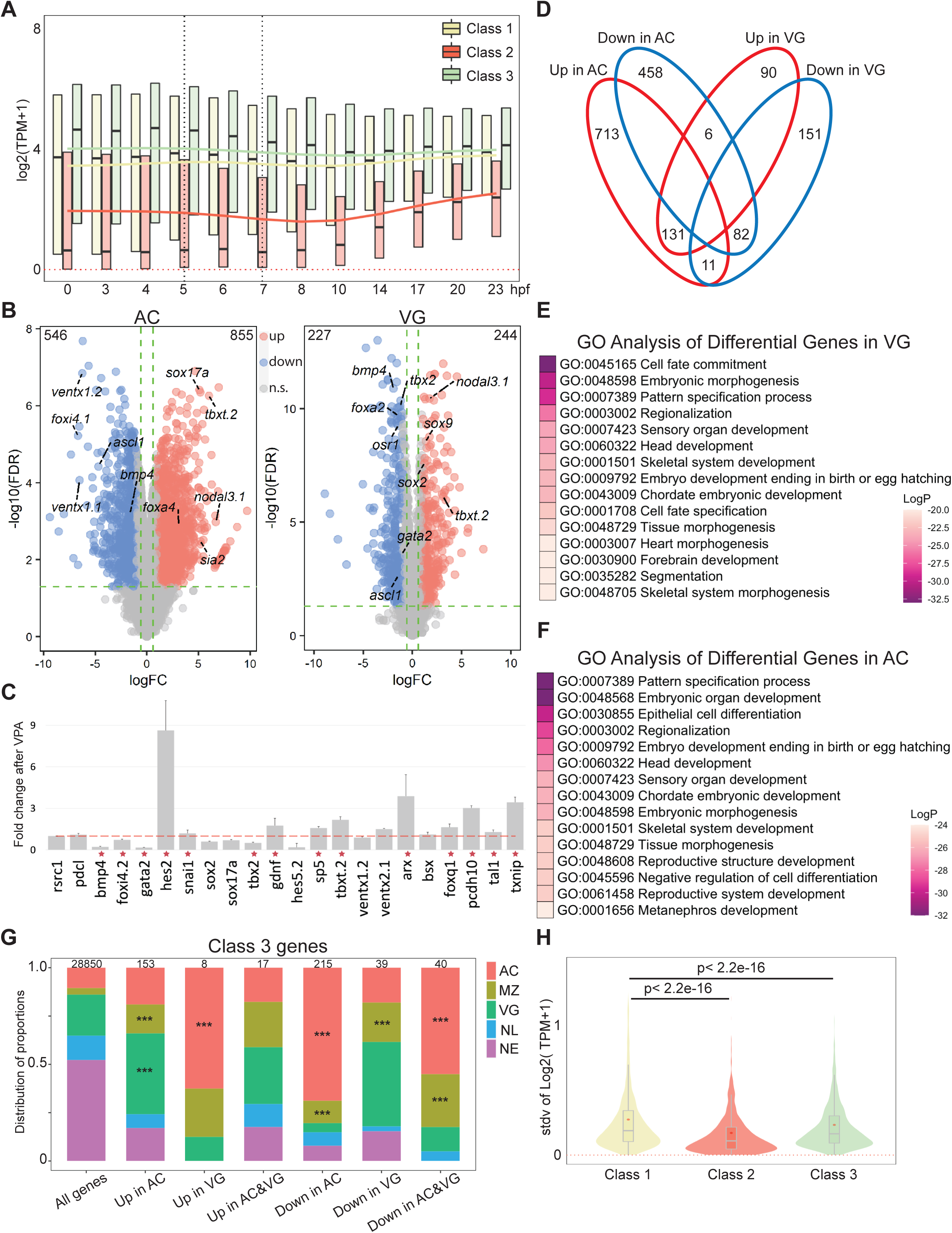
The Integrity of germ layer transcriptomes requires HDAC activity. (**A**) Time-course TPM expression of all genes in Classes 1, 2, and 3 up to 23 hpf (hours post fertilization, tailbud st26). Red dotted line denotes zero. Black dotted vertical lines denote the time window when Hdac1 binding is examined. Trend lines for each class are generated by connecting the mean TPM values at each time point. (**B**) Differentially expressed genes after TSA treatment in either animal cap cells (AC) or vegetal mass cells (VG). The total number of differentially expressed genes in each category is listed on the top two corners. X-axis represents log2 transformed gene expression fold changes after TSA treatment, y-axis represents log10 transformed false discovery rate. n.s.: not significant. (**C**) RT-qPCR analysis examining the effect of VPA on TSA-responsive genes in whole embryos. *rsrc1* and *pdcl* are control genes; *bmp4*, *snai1*, *foxi4.2*, *gata2*, *hes2*, *sox2*, *sox17a*, and *tbx2* are Class 1 genes; *gdnf*, *hes5.2*, *sp5*, *txbt.2*, *ventx1.2*, and *ventx2.1* are Class 2 genes; *arx*, *bsx*, *foxq1*, *pcdh10*, *tal1,* and *txnip* are Class 3 genes. Red stars denote genes regulated in the same direction by both VPA and TSA treatments. The error bars represent the variation from 2 technical replicates. (**D**) Venn diagram comparing the genes differentially regulated by HDAC activity in AC and VG germ layers. (**E, F**) Gene ontology enrichment analyses of genes in (**E**) AC (963 genes) or (**F**) VG (334 genes) that are differentially expressed upon HDAC inhibition. The numbers of genes used for gene ontology enrichment analyses are smaller because the analyses only included genes with matched gene synonym to *Homo sapiens*. (**G**) Spatial expression patterns of Class 3 genes affected by TSA in proportions. The total number of genes in each category is listed at the top of each bar. Only *** denoting *p*< 0.001 (Fisher’s exact test) is shown. AC: animal cap, presumptive ectoderm; MZ: marginal zone, presumptive mesoderm; VG: vegetal mass, presumptive endoderm; NL: non-localized genes; NE: non-expressed genes. Parameters of spatial categories are detailed in Supplemental Methods. (**H**) The standard deviation of TPM expression between five germ layer explants (Blitz et al. 2017) for Class 1, 2, and 3 genes. *p*-values are calculated using Student’s *t*-test.

**Supplemental Table S1:**
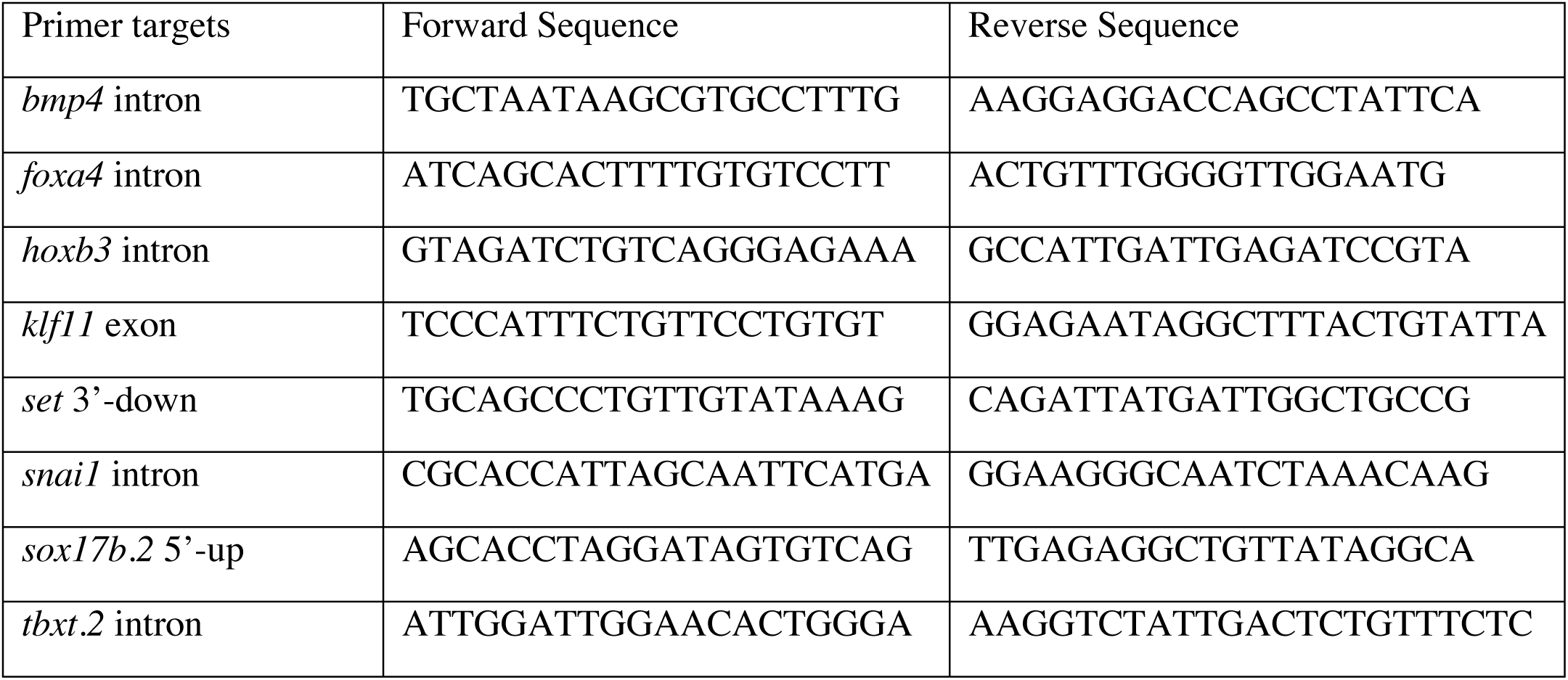
Primer Sequences for sequential ChIP-qPCR

**Supplemental Table S2:** Normalization Factors of ChIP spike-in strategy. Table listing all spike-in normalization factors. Reads mapped to *Xenopus tropicalis* (frog) for all second replicate samples are already **downsampled to 25%**. Red numbers indicate the base ratio line for each column. The ratios for reads mapped to H2Av peaks boxed in yellow are used for all later calculations; reads mapped to *Drosophila melanogaster* (fly) serves as a control to show the highly consistent ratios between whole-genome and H2Av peak mapped reads. The ratios for reads between replicate-merged samples are boxed in blue (corresponding to ratios used in Figure 4F, S4J), which is calculated as sum of Ratio_frog_ *Ratio_fly_ between two replicates. The ratios for reads between explant-merged samples are boxed in magenta (corresponding to ratios used in Figure S4G), which is calculated as sum of Ratio_frog_ *Ratio_fly_ between two replicates followed by two explants.

| Sample | Reads mapped to frog | Ratio (frog) | Reads mapped to fly | Ratio (fly) | Reads mapped to H2Av | Ratio (H2Av) | Ratio of merged reps | Ratio of merged explants |
| --- | --- | --- | --- | --- | --- | --- | --- | --- |
| AC_DMSO_rep1 | 43417738 | 1.17 | 2817756 | 4.09 | 615449 | 3.18 | 18.07 | 33.57 |
| AC_TSA_rep1 | 127763576 | 3.43 | 1175034 | 1.71 | 229326 | 1.18 | 12.49 | 24.66 |
| VG_DMSO_rep1 | 41215108 | 1.11 | 2749022 | 3.99 | 657611 | 3.40 | 15.49 |  |
| VG_TSA_rep1 | 124387262 | 3.34 | 688352 | 1 | 193553 | 1 | 12.16 |  |
| AC_DMSO_rep2 | 47280904 | 1.27 | 10230400 | 14.86 | 2187348 | 11.30 |  |  |
| AC_TSA_rep2 | 86344294 | 2.32 | 3320496 | 4.82 | 703883 | 3.64 |  |  |
| VG_DMSO_rep2 | 37237950 | 1 | 9530640 | 13.85 | 2268171 | 11.72 |  |  |
| VG_TSA_rep2 | 101907192 | 2.74 | 2557466 | 3.72 | 622410 | 3.22 |  |  |

**Supplemental Table S3:**
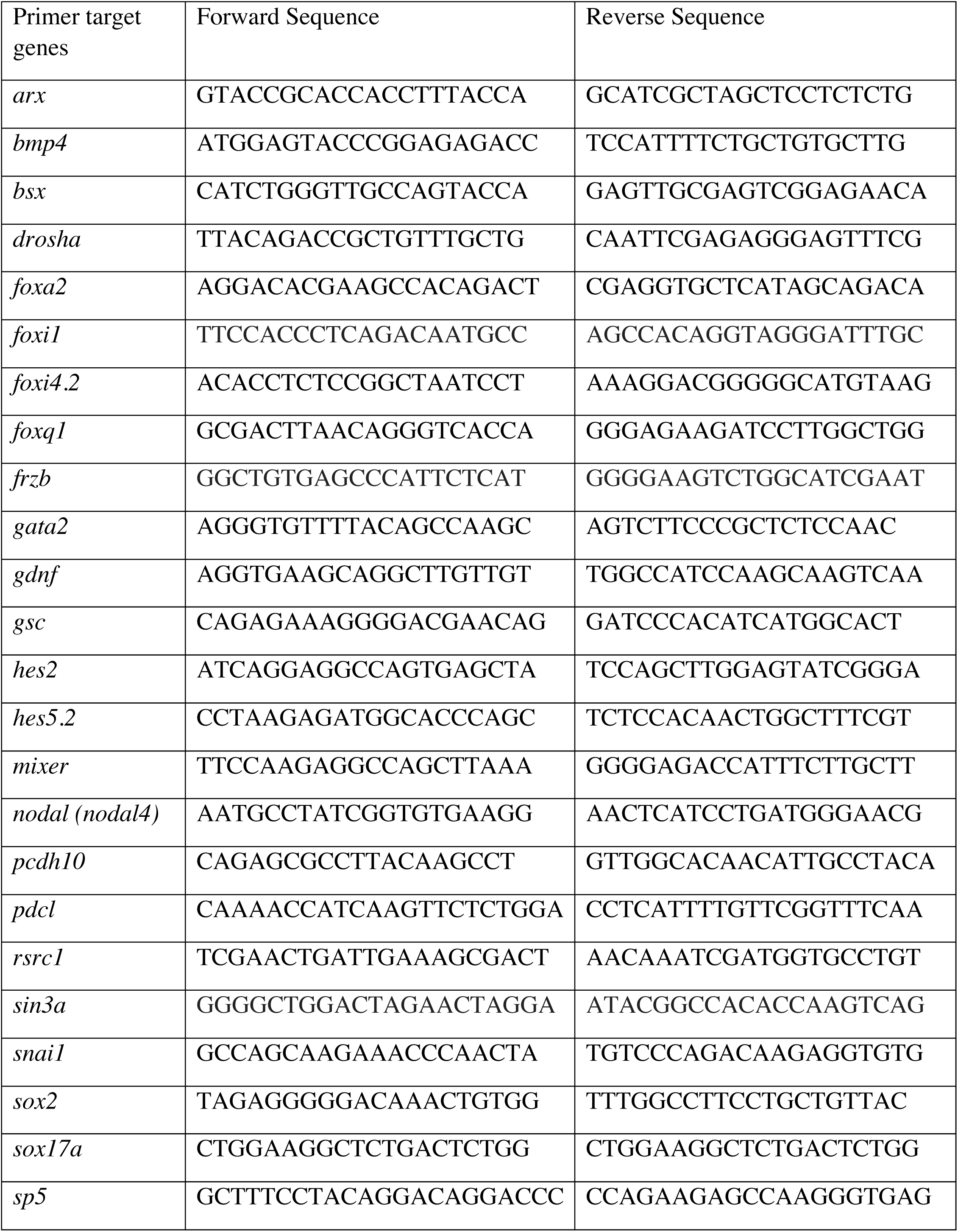
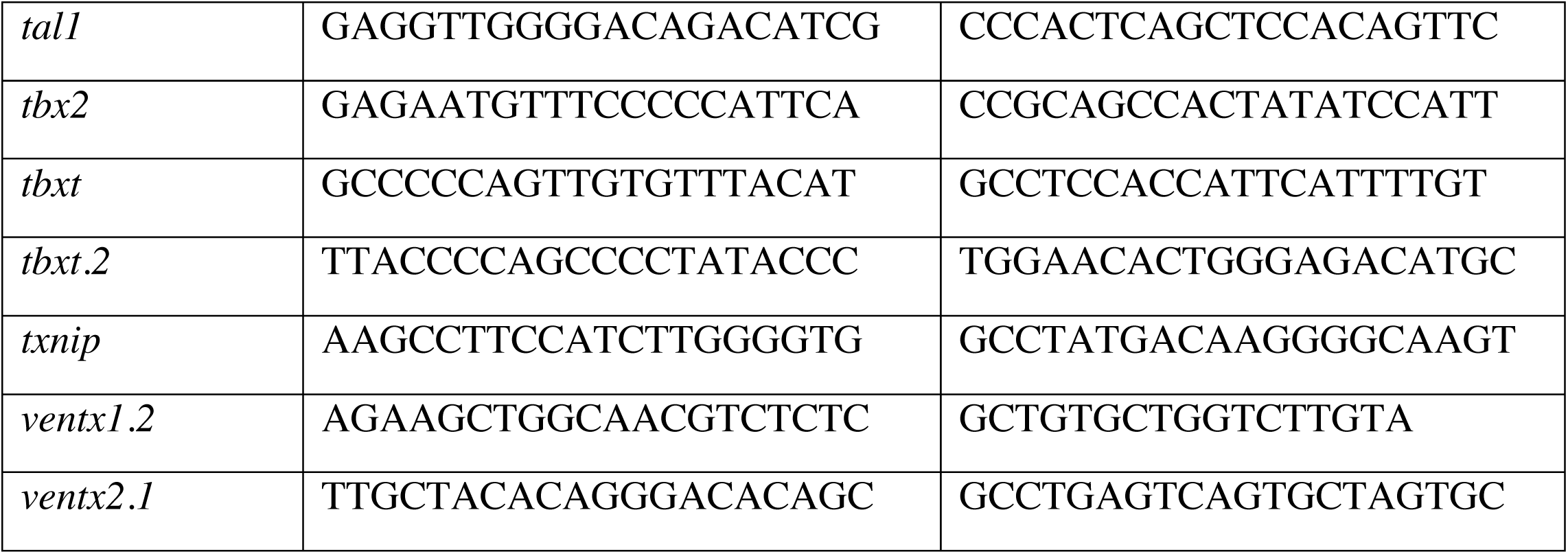
Primer Sequences for RT-qPCR

## Notes

### Competing Interest Statement

The authors have declared no competing interest.

